# HMMRATAC: a Hidden Markov ModeleR for ATAC-seq

**DOI:** 10.1101/306621

**Authors:** Evan D. Tarbell, Tao Liu

## Abstract

ATAC-seq has been widely adopted to identify accessible chromatin regions across the genome. However, current data analysis still utilizes approaches initially designed for ChIP-seq or DNase-seq, without considering the transposase digested DNA fragments that contain additional nucleosome positioning information. We present the first dedicated ATAC-seq analysis tool, a semi-supervised machine learning approach named HMMRATAC. HMMRATAC splits a single ATAC-seq dataset into nucleosome-free and nucleosome-enriched signals, learns the unique chromatin structure around accessible regions, and then predicts accessible regions across the entire genome. We show that HMMRATAC outperforms the popular peak-calling algorithms on published human ATAC-seq datasets. We find that single-end sequenced or size-selected ATAC-seq datasets result in a loss of sensitivity compared to paired-end datasets without size-selection.

## INTRODUCTION

The genomes of all known eukaryotes are packaged into a nucleoprotein complex called chromatin. The nucleosome is the fundamental, repeating unit of chromatin, consisting of approximately 147 base pairs of DNA wrapped around an octet of histone proteins (1). The eviction of nucleosomes into nucleosome-free regions (NFRs) makes DNA more accessible to various DNA binding transcription factors (2). Transcription factor bindings exert spatiotemporal control of gene expression, which is critical in the establishment of cellular identity during development, cellular responses to stimuli, DNA replication and other cellular processes (3). The genome-wide investigations on nucleosome positioning found the interplay between transcription factor binding to nucleosome organization at accessible chromatin where the binding sites are nucleosome-depleted and surrounded by precisely phased nucleosomes (4).

Several assays exist to identify open chromatin regions in a genome-wide manner. These include DNase-seq, which utilizes the DNase I nuclease (5), FAIRE-seq, which utilizes differences in polarity between nucleosome-bound and nucleosome-free DNA (6), and ATAC-seq, which uses a transposase to cut into accessible DNA selectively (7). Although each of these assays identifies some unique open chromatin regions, they are generally highly correlated in their identifications (7). Whereas DNase-seq and FAIRE-seq are complex protocols that require, on average, one million cells, ATAC-seq is a simple three-step protocol, which is optimized for fifty thousand cells and can be performed on as few as 500 cells or at a single cell level (7,8). ATAC-seq has become popular over the years, and the Cistrome Database (9), in an effort to collect all publicly available functional genomics data, has listed nearly 1,500 datasets for human and mouse.

Due to steric hindrance, the Tn5 transposase used in ATAC-seq preferentially inserts into NFRs. However, it is also possible for the transposase to insert into the linker regions between adjacent nucleosomes, resulting in larger (over 150 bps) DNA fragments, which correspond to the integer numbers of adjacent nucleosomes (7). The DNA fragments are constructed in a paired-end library for sequencing, and after mapping both sequenced ends of each fragment to the genome sequence, we can infer their fragment lengths according to the observed mapping locations, or the insertion length. As demonstrated in (7), if we plot the observed fragment length versus frequency, we will see a multi-modal distribution that creates different modes representing transposase insertion into NFRs and linker regions between nearby phased nucleosomes up to +4 nucleosome. This fact allows ATAC-seq to elucidate multiple layers of information relative to the other assays. Although computational tools exist for DNase-seq, FAIRE-seq and ChIP-seq (10), that can be and are used for ATAC-seq analysis, such as MACS2 (11) and F-Seq (12), these would fall short since they only utilize a subset of information, usually the nucleosome-free signals. NucleoATAC (13) was designed to separate an ATAC-seq dataset into its nucleosome-free and nucleosome components then to identify nucleosome positions. However, NucleoATAC only refines a predefined peak set, rather than utilize the nucleosomal information in the peak calling process. To date, there are no dedicated peak-callers specifically to account for both NFR and nucleosomal signals at the same time from ATAC-seq.

We present here HMMRATAC, the Hidden Markov ModeleR for ATAC-seq, a semi-supervised machine learning approach for identifying open chromatin regions from ATAC-seq data. The principle concept of HMMRATAC is built upon “decomposition and integration”, whereby a single ATAC-seq dataset is firstly decomposed into different layers of coverage signals corresponding to the sequenced DNA fragments originated from NFRs or nucleosomal regions; and then the relationships between the layers of signals at open chromatin regions are learned in a Hidden Markov Model and utilized for predicting open chromatins. Our method takes advantage of the unique features of ATAC-seq to identify the chromatin structure more accurately. We found that HMMRATAC was able to identify chromatin architecture and the most likely transcription factor binding sites. Additionally, compared with existing methods used for ATAC-seq analysis, HMMRATAC outperformed them in most tests, including recapitulating active and/or open chromatin regions identified with other assays.

A typical analysis pipeline for ATAC-seq would begin with aligning the sequencing reads to a reference genome using aligner such as BOWTIE2 (14) or BWA (15), then the identification of accessible regions or “peaks” by HMMRATAC, and then downstream analysis such as motif enrichment using MEME (16) or footprint identification in the accessible peaks with CENTIPEDE (17); differential accessibility analysis with Diffbind (18); and association studies with other data sets, such as with gene expression data with BETA (19). Quality control measurements would also take place at each step, such as calculating sequence quality before alignment using FASTQC (20) or after the alignment and performing replicate correlation during peak calling using ATACSeqQC (21). We envision HMMRATAC becoming the principle peak-calling method in such a pipeline.

## METHODS

### Preprocessing of ATAC-seq data

The human GM12878 cell line ATAC-seq paired-end data used in this study was publicly available and downloaded under six SRA (22) accession numbers SRR891269-SRR891274. There are three biological replicates generated using 50,000 cells per replicate and other three generated using 500 cells per replicate. Each dataset was aligned to the hg19 reference genome using BOWTIE2 (14). After alignment, each group of replicates (either 50,000 cells or 500 cells) were merged together, sorted and indexed. Reads that had a mapping quality score below 30 or that were considered duplicates (exact same start and stop position) were removed from the merged files. It should be noted that HMMRATAC will remove duplicate and low mapping quality reads by default, although some other algorithms do not.

The merged, filtered and sorted BAM files, created as described above, were the input for MACS2. HMMRATAC took the sorted paired-end BAM file and its corresponding BAM index file as the main inputs. F-Seq requires a single-end BED file of alignment results as its input. To generate this BED file, we converted the paired-end BAM file into a BED file, using an in-house script, and split each read pair into forward and reverse strand reads.

The human monocyte data from (23) was publicly available and downloaded from the Cistrome Database (9) as aligned BAM files to the hg19 reference genome. The data corresponds to Gene Expression Omnibus accessions GSM2325680, GSM2325681, GSM2325686, GSM2325689, and GSM2325690. One replicate per condition was used and processed in the same way as described above, including merging and filtering on the BAM files.

### The HMMRATAC algorithm

The HMMRATAC algorithm is built upon the idea of “decomposition and integration”, and based on the observation of distinct nucleosome organization at accessible chromatin (2–4,7) (see **Results, Figure 1A**, and **Supplemental Figure S1**). A single ATAC-seq dataset is firstly decomposed into different layers of coverage signals corresponding to the sequenced DNA fragments originated from NFRs or nucleosomal regions; and then the relationships between the layers of signals at open chromatin regions are learned in a Hidden Markov Model and utilized for predicting accessible regions across the whole genome (**Figure 1B**). The detailed steps of HMMRATAC algorithm are described as follows.

**Figure 1.**
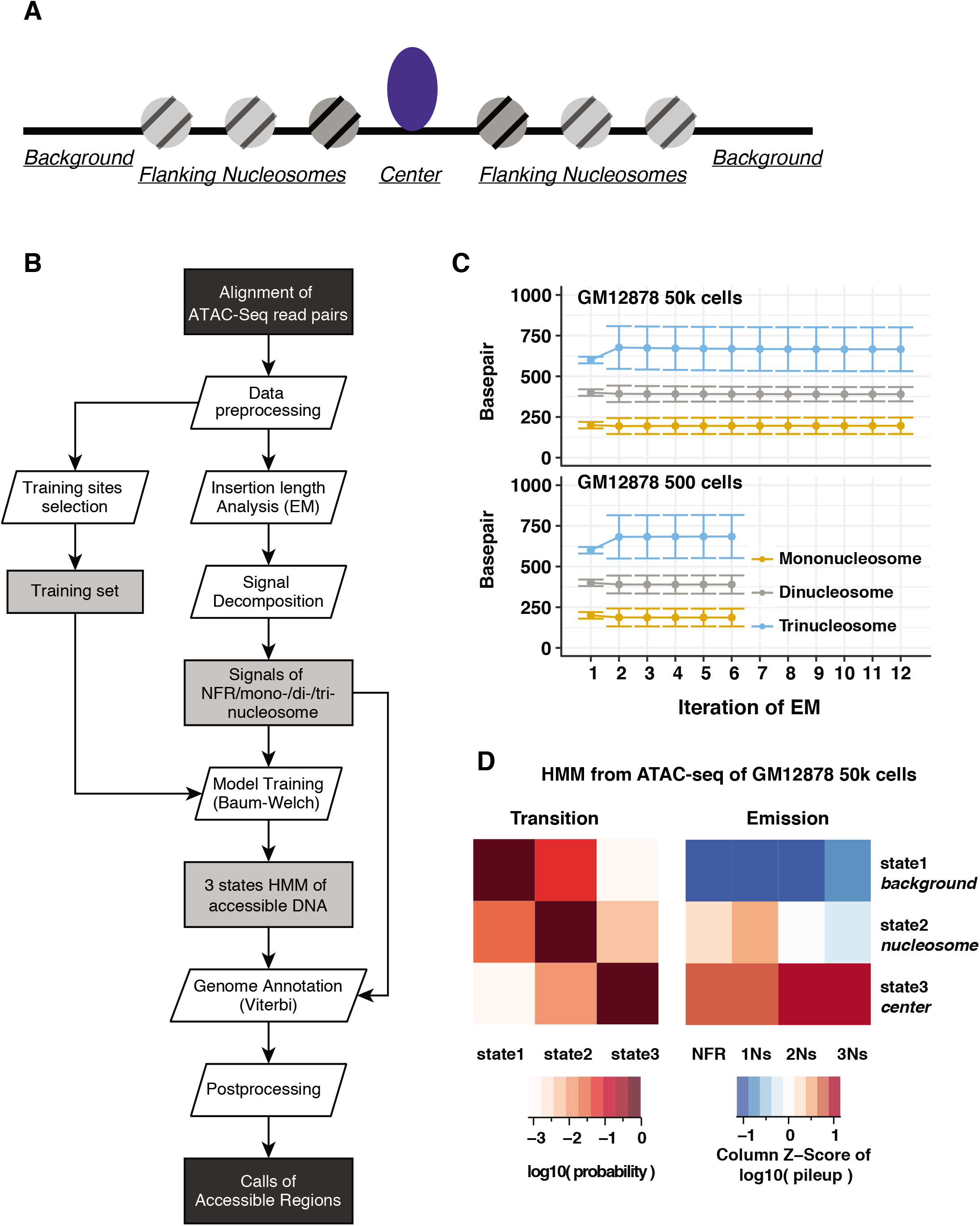
The HMMRATAC Algorithm. **(A)** A schematic to represent the structure of an accessible region. **(B)** The algorithmic workflow. **(C)** Estimation of fragment length parameters through the Expectation-Maximization algorithm. Mean and standard deviation parameters per EM iteration for three nucleosomal distributions are consistent among datasets using 50,000 and 500 cells. The parameters for NFR signals are fixed during EM training so they were not plotted. Iteration #1 shows the initial parameters. EM was applied to pooled biological replicates. The parameters per biological replicate were shown in **Supplemental Figure S3. (D)** The transition and emission parameters of the Hidden Markov Model trained through the Baum-Welch algorithm on the 50,000 GM12878 cells ATAC-seq data.

#### Segregation of ATAC-seq signals

After the preprocessing step that eliminates duplicate reads and low mapping quality reads, the main HMMRATAC pipeline (**Figure 1B**) begins by separating the ATAC-seq signal into four components, each representing a unique feature. These features are nucleosome-free regions (NFRs), which are most likely to occur within the open chromatin region itself, and three nucleosomal features, representing mono-, di- and tri-nucleosomes (1Ns, 2Ns, and 3Ns), respectively. We utilize four distributions to represent the four signal tracks: an exponential distribution (*P_s_*(*β|x*)) for the nucleosome-free track and three Gaussian distributions (*P_m_*(*μ_m_, σ_m_*), *P_d_*(*μ_d_, σ_d_|x*), *P_t_*(*μ_t_, σ_t_|x*)) to represent the mono-, di- and tri-nucleosomal signal tracks, where *x* represents the observed insertion length of a reads pair, *β, μ_m_, μ_d_*, and *μ_t_* represent the means, *σ_m_, σ_d_* and *σ_t_* represent the standard deviations of the corresponding distribution.

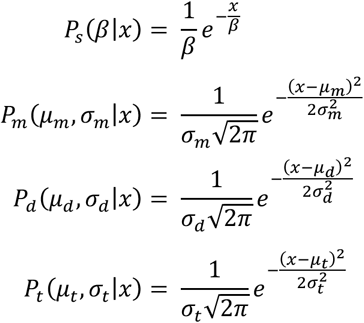

These distributions were utilized by Buenrostro et al. to identify their cutoffs for fragment separations (7). Although sequence read data is discrete, the fact that the expectation value of each Gaussian is large (>100 bps) and the number of observations is high, the discrete distribution, such as a Poison distribution, can be approximated as a Gaussian. The mean value of the exponential distribution for NFRs is set as a fixed value at runtime, either as the default value or a user-defined value. The parameters of three Gaussian distributions are initialized at runtime with default or user-defined values and then updated using the expectation maximization (EM) algorithm for Gaussian mixture models. These initial values were the same as those used by Buenrostro et al. (7) to define their distributions for the chromatin features. Briefly, each nucleosomal fragment, those larger than 100 bps, is classified as belonging to one of the nucleosomal distributions based on its weighted probability. Once each fragment has been classified, new means, standard deviations, and weights are calculated for each distribution and become the new values for the mixture model. This process continues iteratively until the change in the mean values between iterations is less than a reasonable epsilon value. At this point, the model is assumed to have reached convergence, and the EM process is halted (**Figure 1C**). In order to decrease the time required for the EM algorithm, HMMRATAC randomly sub-selects 10% of all the fragments to use as the training data. Once the parameters of the four distributions have been determined, HMMRATAC creates four genome-wide signal tracks. For every genomic position, all of the fragments that occupy that position are determined. Each track is then incremented by the probability calculated in the above equation that a particular fragment belongs to the tracks’ corresponding distribution. We then use a square root transformation to get the data into continuous space. Although the transformations are commonly done with logarithmic transformations, we chose square root because our data is all positive values but also contains zeroes.

#### Training the Hidden Markov Model of accessible chromosomal regions, and refining the model with the Baum-Welch algorithm

Once the four signal tracks have been created, the main HMMRATAC process begins. HMMRATAC first identifies up to 1000 training regions throughout the genome, with which it learns the model. These training regions are determined by scanning the genome for regions whose read coverage fold changes over background falls within a certain range. The default lower and upper limit of fold changes, as used in the Result section, is set between 10 and 20.

These regions are then extended by 5 kbps in either direction, in order to ensure that background regions are included in the training set. It is also possible for users to provide a BED file containing predefined training regions. The initial HMM has proportional initial probabilities, which are not updated during training, as well as proportional transition probabilities, which are updated during training. The emission probabilities are calculated with k-means clustering, by separating the training regions into 3 clusters with random initialization and calculating the means and covariance matrices for the four signal tracks for each cluster. The four signal tracks, across the training regions, are the only input to the k-means algorithm. We model the emissions as a multivariate Gaussian distribution since this assumption has been successfully adopted in other HMM-based genome segmentation methods such as hiHMM (24,25) and Segway (26,27). The three clusters represent the three states of our model, corresponding to the *center* of the open region with overall high enrichment of all of the four types of signals, the *nucleosomes* with low NFR signal but moderate nucleosomal signals, and a *background* state with low enrichment of any signals, respectively. It is also possible to create a model using a different number of states instead of 3, although this option is not generally recommended. Once this initial HMM is created, the transition and emission parameters are updated with the Baum-Welch algorithm (28) until convergence (**Figure 1D**). Furthermore, to demonstrate the robustness of the training approach, we ran HMMRATAC three times on the same GM12878 ATAC-seq data with different random initializations for k-means and found that the k-means centers and HMM parameters were virtually identical, as shown in **Supplemental Table S1**.

#### Annotating with the Viterbi algorithm

Once the HMM has been created and refined, the Viterbi algorithm (29) is used to classify every genomic position into one of the three states of our model. In our experience, when Viterbi encountered a high coverage region, it was interrupted and mistakenly called the entire regions following the high coverage region as either a peak or a background region. Therefore, regions whose z-scored coverage is above a certain cutoff, either a default or user-defined value, are masked prior to the Viterbi algorithm. Additionally, it is possible to include a list of genomic regions to mask along with the extremely high coverage regions, such as annotated blacklisted sites provided by the ENCODE project. We have found that 91% of the high coverage regions that are masked from the Viterbi algorithm in the GM12878 cells are annotated blacklist regions (downloaded from (30)). A genome-wide BedGraph of all state annotations can be reported by HMMRATAC, although this is not the default behavior. By default, HMMRATAC will take all regions classified as belonging to the *center* state and whose length is above a default or user-defined threshold and merge them with the *nucleosome* states located both upstream and downstream of the *center* state. We generally consider the nearby nucleosomes to be part of the regulatory region (See **Results**), and therefore the region is then reported as a peak in gappedPeak format (31). Each reported region is given a score, which represents the maximum read coverage of the *center* state. Additionally, HMMRATAC also supplies a file containing the summits of each region. To calculate a summit, HMMRATAC first filters the read coverage signal throughout the *center* state using a Gaussian smoothing function with a window size of 120 bps. It then calculates the position that has the maximum read coverage and reports that position as the summit.

#### Software implementation

HMMRATAC software is implemented using JAVA language. It should be noted that at all stages of the algorithm, HMMRATAC does not store the read or coverage information in memory. Any information needed by the program is read from the desired file and used at the specific time. This was incorporated to reduce the memory usage of HMMRATAC. However, because of this feature, it is not recommended to run HMMRATAC simultaneously with the same data files. It is, however, possible to process multiple different data sets in parallel. In terms of system requirements, the only required component is to have Java 7 or higher installed. We have tested HMMRATAC on OpenJDK 7, standard edition JRE 8 and the standard edition JRE 10. In terms of hardware requirement, although all results involved in this study were generated on our server equipped with AMD Opteron™ Processor 6378 and 500GB memory, HMMRATAC can successfully process human ATAC-seq datasets on modest personal computers such as the author’s laptop equipped with a 2.8GHz Intel Core i7 processor and 16 GB of DDR3 memory. The actual maximum memory usage and the runtime of HMMRATAC mainly depend on the parameter settings. As an example, when the window size for Viterbi decoding (option --window) was set to 2,500 kbps, the maximum memory usage decreased from 24GB, when using the default window size of 25,000 kbps, down to 13 GB, on the same merged GM12878 ATAC-seq dataset. We recognize that HMMRATAC takes longer than most algorithms and are currently optimizing it in order to decrease runtime. We will also introduce a parallel computing ability, which we believe will dramatically reduce the runtime. These efforts are active and ongoing and will be incorporated into future releases.

### Evaluation

#### Calls from MACS2

MACS version 2.1 was used to call peaks using the sorted paired-end BAM files created as described above. With each run the input file format (option -f) was set to “BAMPE” and the option for keeping duplicate reads (option --keep-dup) was set to “all”, as duplicate reads had already been discarded in pre-processing. MACS2 was run with the local lambda option turned off (option --nolambda) and was run with q-value cutoffs (option -q) set to 0.5, 0.1, 0.05 (default), 0.005, 0.0005 and 0.00005. It was also run with a p-value cutoff (option -p) set to 0.6 and 0.3.

#### Calls from F-Seq

F-Seq version 1.84 was used to call peaks using the single-end BED files created as described previously. With each run, the output was reported as a BED file (option -of). The standard deviation cutoff (option -t) was set to 0.1, 1, 4 (default), 6, 7, 8, 10, 15 and 20.

#### Calls from HMMRATAC

HMMRATAC version 1.2 was used to call peaks using the sorted BAM file. The upper limit fold-change for choosing training regions (option -u) was set to 20 with corresponding lower limits (option -l) set to 10. These settings are the HMMRATAC defaults. The blacklisted sites for hg19 were provided, which were masked from the program and all subsequent steps. For the precision, recall and false positive rate calculations, the HMMRATAC peaks were filtered by minimum score cutoffs of 0, 5, 10, 15, 20, 25, 30, 35, 40 and 45. Additionally, for two monocyte samples (accession numbers GSM2325689 – RPMI_4h and GSM2325681 – BG_d1) HMMRATAC did not produce a typical model with the default settings (-u 20 –l 10). The ideal model should show the highest NFR, 1Ns, 2Ns and 3Ns signals in the third (center) state and the second highest values of all four signals in the second (nucleosome) state. In this case, we found that the second state (nucleosome) had the highest NFR signal while the third (center) state had the highest 1Ns, 2Ns, and 3Ns signals. Re-running it with more stringent parameters (-u 40 –l 20) solved that problem and produced a model that had the highest signal values in the third state and the second highest in the second state. These peak results from HMMRATAC were then filtered by the same cutoffs as the other samples and tested in the same way. All the transition and emission parameters learned by HMMRATAC from GM12878 50k cells, 500 cells, five human monocyte samples are summarized in **Supplemental Table S2**.

#### Compiling the “Gold Standard” lists and “True Negative” set

We tested the performance of HMMRATAC and other methods on ATAC-seq datasets of GM12878 cells against three proxies of “gold standard” datasets. The first one, which we call “active regions”, is made by combining two chromatin states “Active Promoters” and “Strong Enhancers” generated in GM12878 cells using chromHMM (32,33) (downloaded from UCSC genome browser track wgEncodeBroadHmmGm12878HMM). The other two sets were DNaseI hypersensitive sites (UCSC track wgEncodeAwgDnaseUwdukeGm12878UniPk) and FAIRE accessible regions (wgEncodeOpenChromFaireGm12878Pk) (34) called from DNase-seq and FAIRE-seq on the GM12878 cell line. Because the DNase-seq and FAIRE-seq regions were called by the ENCODE project (35) from traditional methods including Hotspot (36) and F-Seq (12), the evaluation based on either of them is less unbiased compared with the test based on chromatin states which integrate multiple independent experiments. We found that there were numerous cases throughout the genome where clusters of DNaseI or FAIRE regions were in close proximity to one another. These clusters tended to have broad enrichment in either DNase-seq or FAIRE-seq signals. Therefore, we merged all DNaseI or FAIRE regions that were within 2 kbps of each other into a single region and used these merged sets as the proxies to the gold standard.

The “real negative” set was made with the “Heterochromatin” state annotation from chromHMM. When using the “active region” set as “gold standard”, no modifications were necessary because the three states are mutually exclusive. When using either the DNaseI or FAIRE sets, any “heterochromatin” states that overlapped (by any amount) a DNaseI or FAIRE region were excluded from the final “real negative” set. Additionally, any DNaseI or FAIRE peak that overlapped any “heterochromatin” state (by any amount) was, likewise, excluded from its respective “gold standard” set.

The “gold standard” set used for the human monocyte ATAC-seq data were compiled as follows. For each condition, 2 replicates of H3K27ac, H3K4me1 and H3K4me3 ChIP-seq datasets, one replicate of H3K9me3 from the same study (23) were downloaded from the Cistrome Database (9) as aligned BAM files. Each file was processed by MACS2 with the p-value parameter (-p option) set to 0.15. We set this loose cutoff, so that, when combining active and heterochromatin marks, the true and false sets can be more exclusive. In order to identify a loose cutoff, we performed MACS2 cutoff-analysis (callpeak --cutoff-analysis) approach to identify the critical cutoff below which the genomics background noises start to be picked up by peak caller. The resulting peaks were sorted by the −log10(FDR) value and the top 10000 peaks for each sample were kept. The “active regions” true set was created as the genomic regions overlapped with either H3K27ac, H3K4me1 or H3K4me3 but not overlapped with H3K9me3 peaks. This list of “active regions” became the proxy of “gold standard” set for each condition. Any H3K9me3 peak that did not overlap the “active region” set was kept as the “real negative” set for that condition. The accession numbers for all above histone modification ChIP-seq data are listed in **Supplemental Table S3**.

#### Calculating Precision, Recall, False Positive Rate and Area Under the Curves

Let Predicted Positive (PP) represent the number of base pairs identified as accessible regions from any method. Let Real Positives (RP) represent the number of base pairs in the “gold standard” sets. Let Real Negatives (RN) represent the number of base pairs in the “real negative” set. Let True Positive (TP) represent the number of overlapping base pairs between a called peak and a real positive. Let False Positive (FP) represent the number of overlapping base pairs between a called peak and a real negative.

Then the *precision* or positive predictive value (PPV) is calculated as PPV = TP/(TP+FP). The *Recall*, or true positive rate (TPR), is calculated as TPR = TP/RP. The *false positive rate* (FPR) is calculated as FPR = FP/RN. The number of overlapping base pairs was calculated with the BEDTools intersect tool (37). After the precisions, recalls and false positive rates were calculated for accessible regions predicted by a given method under various cutoff values, the approximate area under the curve (AUC) was calculated for the *recall* vs. *false positive rate* (ROC) curves and for the *precision-recall* (PR) curves, after adding extreme points to the unreachable ends of the curves. The extreme points to the ROC curves are TPR=0 and FPR=0, and TPR=1 and FPR=1; the extreme points to the PR curves are TPR=0 and PPV=1, and TPR=1 and PPV=L_true_/(L_true_ + L_false_) where the L_true_ is the total length of real positive set and the L_false_ is the total length of the real negative set.

## RESULTS

### The distinctive pattern of ATAC-seq fragments around known accessible elements

In order to investigate the ATAC-seq signal profile around accessible genomic regions, we took the ATAC-seq data in the human GM12878 cell line from Buenrostro et al. (7) and separated the fragments into short nucleosome-free (under 100 bps) and mono-nucleosome (180-250 bps) fragments based on cutoffs identified by Buenrostro et al. These cutoffs were determined by fitting the fragment length distribution to several simulated distributions modeling nucleosome and nucleosome-free transposition frequencies (7). We plotted the two types of fragments around the centers of DNaseI hypersensitive sites (DHSs) in the same cell line, identified by ENCODE (35) (**Supplemental Figure S1**). We observed a clear, symmetrical pattern around the centers of these sites characterized by an enrichment of the nucleosome-free fragments and flanking enrichment of the mono-nucleosome fragments. We were able to visually discern at least three distinct regions around an open site: 1) the *center*, characterized by an enrichment of nucleosome-free fragments and flanked by nucleosome fragments; 2) the *nucleosome regions*, characterized by an enrichment of nucleosome fragments; 3) and the *background*, characterized by a depletion in any fragments (**Figure 1A; Supplemental Figure S1**). Based on this observation and existing knowledge on the nucleosome positioning around the accessible chromatin (2–4), we hypothesized that such pattern would exist at the open chromatin throughout the genome and that by learning and recognizing this pattern computationally in a single ATAC-seq dataset, we would identify the open chromatin regions with higher confidence than any other existing method.

### Signal decomposition through the probabilistic approach

In the first ATAC-seq paper by Buenrostro et al., the authors separated the fragments according to their insertion lengths into four populations representing those from the NFRs and those spanning 1, 2 or 3 nucleosomes (7). Non-overlapping and non-adjacent cutoffs were used. For example, lengths below 100 bps represent nucleosome-free signals and lengths between 180 and 250 bps represents mono-nucleosome signals. We consider such cutoff-based strategy not a general practice for the following two reasons. First, different species or different cell lines may have different nucleosome spacing and these differences may not be adequately understood (38,39). Secondly, since the fragments whose size falls between the non-overlapping and non-adjacent cutoffs are discarded (e.g. those between 100 and 180 bps long), the recall (or sensitivity) to identify accessible regions will be reduced (**Supplemental Figure S2**). As a fact, about 15% of the entire data generated from the original paper are ignored with their strategy. We addressed the above two challenges in a probabilistic approach in HMMRATAC.

First, we utilize the Expectation-Maximization (EM) algorithm to identify the optimal parameters for the four different distributions of fragment lengths. Using the distributional assumptions of Buenrostro et al., we create four distributions to represent four different chromatin features: nucleosome-free regions (NFRs), mono-nucleosomes (1Ns), di-nucleosomes (2Ns) and tri-nucleosomes (3Ns). HMMRATAC uses a mixture model and Expectation-Maximization (EM) algorithm to identify optimum parameters. As an example, HMMRATAC identified the optimum fragment lengths representing 1Ns, 2Ns, and 3Ns from ATAC-seq data in the human GM12878 cell line (7), generated by merging three replicates, as being 195 bps, 396 bps, and 693 bps, respectively (**Figure 1C**). As a demonstration of different nucleosome spacing in different datasets, HMMRATAC determined the optimum fragment lengths for the 1Ns, 2Ns, and 3Ns from ATAC-seq data in human monocytes treated with RPMI for one day (23) as 186 bps, 368 bps, and 549 bps, respectively. Additionally, the values for any of the individual replicates, in the GM12878 data, were never more than 5 bps off from the average values listed above for the merged datasets (**Supplemental Figure S3**). Not all ATAC-seq datasets show a clear multi-modal pattern, especially datasets from fewer cells. As an example, the fragment distribution generated by Buenrostro and et al. utilizing 500 cells (GM12878 cells) per-replicate, did not show as clear of a distribution as 50,000 cells per replicate (**Supplemental Figure S4A and S4B**). However, we found our approach can robustly identify the parameters even if the multi-modal pattern is not clear. As we can see from **Figure 1C**, the mean and standard deviation values converged after no more than 12 iterations and the parameters converged to very similar values for either the 50,000-cell per-replicate or 500 cell per-replicate datasets.

Secondly, we propose in HMMRATAC a probabilistic approach to make use of data from the entire spectrum of fragment lengths. We generate four distinct signal profiles in 10 bps resolution, for NFRs, 1Ns, 2Ns, and 3Ns, allowing each mapped fragment to contribute to all profiles with different weights corresponding to its particular fragment length. For each genomic location that is occupied by a particular fragment, all four signal profiles are incremented by the probabilities that this fragment, assigned with an observed fragment length through read mapping, belongs to each of the four corresponding distributions. We tested the performance of a regular HMMRATAC approach with this probabilistic method utilizing the entire data and an alternative approach utilizing only the subset of data satisfying the cutoffs proposed in Buenrostro et al. The two approaches were compared in terms of precision and recall against the “active regions” true set (**Supplemental Figure S2**). We found that our probabilistic method had considerably better recall and precision compared to the cutoff based method. This result proved that the traditional way to discard reads between cutoffs of fragment sizes would hurt the prediction power.

HMMRATAC utilizes the length of an insertion fragment, identified by paired-end sequencing, as an essential and critical piece of information in its processes. As such, HMMRATAC can only be used in paired-end sequencing libraries. Because we observed a decrease in the performance of our method when we used the cutoff based approach, which resulted in the removal of ~15% of all sequencing reads, we wanted to understand the effects of size-selection and paired-end sequencing on ATAC-seq peak calling. We performed two *in silico* size-selections on the data from Buenrostro et al. The first restricted the fragments to those whose length was under 100 bps and the second restricted the fragments to those with lengths under 300 bps. We also treated the data as single-end, by unpairing the read mate pairs. We then used MACS2 (11) to call peaks with each data set, including the original unaltered data and compared the performance to predict the accessible chromatin (**Supplemental Figure S5**). As a result, the original unaltered data performed the best in terms of precision and recall. This indicates that an ATAC-seq assay should better be performed without a size selection step and the resulting fragments should undergo paired-end sequencing, regardless of the peak calling algorithm that is used.

### Integration through Hidden Markov Model

After removing reads with low mapping quality, and masking blacklist regions from downstream analysis, HMMRATAC identifies a set of training regions throughout the genome to train a 3-states Hidden Markov Model (HMM) with multivariate Gaussian emissions, representing the underlying three chromatin states of 1) the *center* of open chromatin, 2) the *nucleosome regions* and 3) the *background* (**Figure 1A**). By default, training regions can be either specified by users or automatically selected where the signal fold-change above background falls within a predetermined range. After signal decomposition, a matrix of 10 bps resolution signals for NFRs, 1Ns, 2Ns, and 3Ns in training regions is used to train the parameters in HMM through the Baum-Welch algorithm (28). The initial state transition probabilities are set equally as 0.33 and the emission parameters for each state are calculated after clustering all locations in the training regions into 3 classes (see **Methods** for detail). **Figure 1D** shows the final parameters for the HMM on ATAC-seq in the GM12878 cell line, by merging three replicates from 50k cells (7). Although the parameters will change depending on the choice of training regions and the datasets provided, several key characteristics are evident in the model and remain mostly stable with different settings and data. State 3 in the HMM model, which HMMRATAC annotated as the *center* state, showed the highest average signal across the four signal profiles. State 2, annotated as the *nucleosome* state, showed the second highest average values. Additionally, as we hypothesized, the results show the *center* state mainly transits from or to the *nucleosome* state.

After learning the model, HMMRATAC takes a matrix of genome-wide NFRs, 1Ns, 2Ns, and 3Ns signals and labels each genomic location with one of the states using the Viterbi algorithm (29). On the 50K cells ATAC-seq data in GM12878 cell line, there were about 4.3 million regions annotated as *background* with an average size of 288 bps (~35% of the human genome), about 4.5 million regions annotated as *nucleosomes* with an average size of 382 bps (~49% of the human genome), and about 226 thousand regions annotated as *center regions* with an average size of 504 bps (~3% of human genome). HMMRATAC finally connected the *center regions* with their adjacent *nucleosomes* and called a total of 87,440 accessible regions with an average size of 1,831 bps (5% of the human genome), excluding those with abnormally high coverage and those below a minimum length. It should be noted that many of the *nucleosomes* are not adjacent to the *center* state. These may represent nucleosomes away from regulatory elements that ATAC-seq is able to identify or may be artifacts due to increased transposition frequency, an issue which can be caught by quality assessment tools such as ATACseqQC (21).

### Recapitulating chromatin architecture at functional elements

We hypothesized that HMMRATAC could identify chromatin architecture around accessible regions, or ideally, nucleosomes near the regulatory elements. To prove this, we explored the distribution of histone modification ChIP-seq signal around our identified accessible regions. We downloaded and analyzed datasets released by ENCODE in the GM12878 cell line (35), including ChIP-seq for CTCF, active histone marks such as H3K4me1, H3K4me3, H3K9ac, and H3K27ac, and repressive histone marks such as H3K9me3 and H3K27me3, as well as sequencing of micrococcal nuclease digestion (MNase-seq). **Figure 2A** shows two loci, one on chromosome 8 that was identified as an accessible region by MACS2 and F-Seq but not HMMRATAC and another locus on chromosome 2 that was identified as accessible by all three methods. At the chromosome 2 locus, we observed that the *center* state was located in regions characterized by depletion in both active histone marks and MNase digestion. The surrounding *nucleosome* states showed enrichment of active histone marks and MNase digestion. We next plotted the same ChIP-seq, DNase-seq and MNase-seq data around all distal (> 10 kbps from any gene promoter) *center* states and used k-means clustering to group the resulting profiles (**Figure 2B**). We found that HMMRATAC states can identify, what appears to be, different chromatin features at the accessible genomic regions. For example, the first cluster (C1) characterized by strong enrichment of H3K27ac relative to H3K4me1 seems to indicate active enhancers while the third cluster (C3) marked by high levels of H3K4me1 and more moderate levels of H3K27ac would suggest weak or poised enhancers (**Figure 2B**). The second cluster (C2), characterized by strong enrichment of CTCF binding, lower levels of H3K4me1, H3K4me3 and acetylation histone marks and a higher level of H3K27me3, seems to suggest that this cluster represents insulators or domain boundaries. The fourth cluster (C4), similar to C2, suggested accessible regions also exist in the DNA elements with repressive functions in euchromatin, potentially bound by factors rather than CTCF. To summarize, these results indicate that HMMRATAC recapitulates the histone architecture around accessible regions.

**Figure 2.**
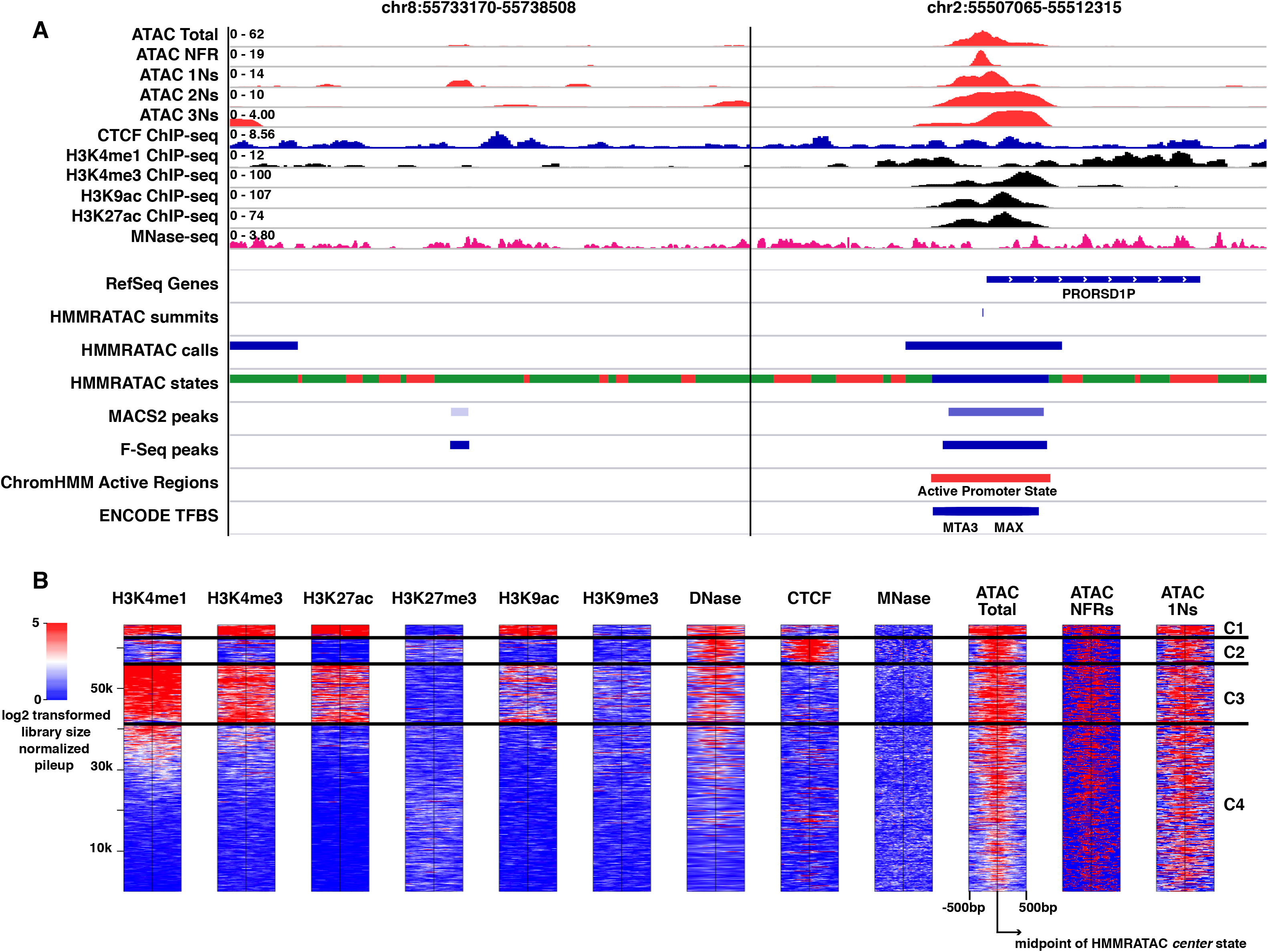
Chromatin Architecture at HMMRATAC Peaks. **(A)** IGV browser snapshot of various signals around a negative locus on chromosome 8 and a positive locus on chromosome 2. Top four tracks show ATAC-seq signals, after being separated by size, the fifth track shows CTCF ChIP-seq signal, the sixth through ninth tracks show various histone modification ChIP-seq signal and the tenth track shows MNase-seq signal. The bottom panel shows RefSeq genes, HMMRATAC calls and state annotations (called with default settings, red: *background* state, green: *nucleosome* state, and blue: *center* state), MACS2 and F-Seq peaks (called with default parameters), the active regions combined from “Active Promoters” and “Strong Enhancers” states from chromHMM (see **Methods**). **(B)** ChIP-seq of H3K4me1, H3K4me3, H3K27ac, H3K27me3, H3K9ac, and H3K9me3, DNase-seq and MNase-seq signals (log2 transformed library size normalized pileup) plotted across all distal (> 10 kbps from gene promoters) *center* states. Data were clustered using k-means algorithm into 4 clusters.

The chromosome 8 locus exemplifies some of the advantages of HMMRATAC over other peak callers. Both MACS2 and F-Seq identified this locus as being accessible, despite the lack of other corroborating evidence, such as transcription factor binding and the presence of active chromatin states. It appears that these methods called the site because of an enrichment of the mono-nucleosome signal. Because neither MACS2 or F-Seq integrate the various chromatin features present in ATAC-seq data and because they are based on signal enrichment only, as opposed to HMMRATAC which incorporates the structure of the element, they both falsely identified this region as accessible, while HMMRATAC identified the region as being a *nucleosome* state.

In order to understand the chromatin features that are enriched in each state across the genome, as the unique result HMMRATAC concluded from a single ATAC-seq data, we plotted the overlap of the 3 hidden states with chromHMM states and histone mark calls from ChIP-seq in the GM12878 cell line based on ENCODE data (**Figure 3A; Supplemental Table S4**). We found that the chromatin states “Active Promoters”, “Strong Enhancer”, “Insulator”, “Poised Promoter” and “Weak Promoter” states mainly overlap with our *center* state, and the “Heterochromatin”, “Repetitive”, “Repressed”, “Transcription Elongation”, and “Weak Transcription” barely overlaps with the *center* state. Interestingly, the “Strong Enhancer” state had a similar amount of overlap with the *nucleosome* state as the *center* state. Combined with our observation that our *nucleosome* state showed elevated levels of active histone marks near accessible regions, we hypothesized that this state may represent positioned nucleosome nearby regulatory elements, those marked by active histone modifications and adjacent to accessible regions. We plotted the H3K4me1 active histone mark around the centers of our *center* states as well as around the centers of the adjacent *nucleosome* states, those directly flanking our *center* states (**Figure 3B**). As we can see, the *center* states show a bimodal enrichment for the histone mark, consistent with MACS2 and F-Seq peaks. Additionally, our adjacent *nucleosome* states show central enrichment for the histone mark. This indicates that the *nucleosome* state, at least those adjacent to our *center* states, are likely nucleosomes close to regulatory elements. As a result, HMMRATAC merges the *center* states with their adjacent *nucleosome* states, both upstream and downstream, to identify the regulatory region as the output.

**Figure 3.**
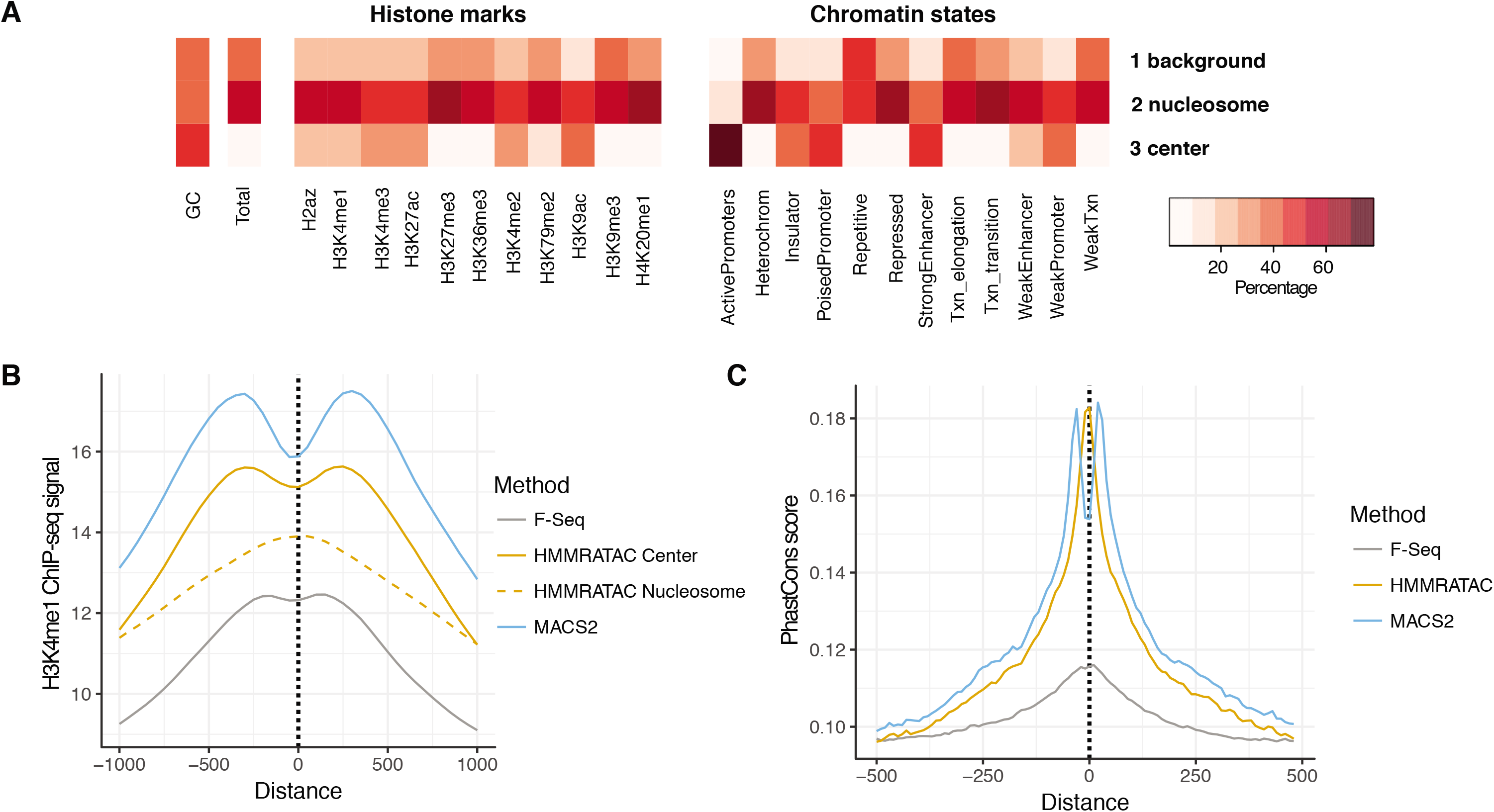
Chromatin features of HMMRATAC states. **(A)** The overlap between either histone modification ChIP-seq peaks or chromatin states in GM12878 and three HMMRATAC states – *background, nucleosome*, and *center*. The color shows the percentage of GC, the percentage of total genome length, or the percentage of histone peaks or chromatin states overlapping with each of HMMRATAC states. Overlaps were calculated in basepair level. **(B)** Distribution of GM12878 H3K4me1 ChIP-seq signal in 1000 bps window surrounding the summits of F-Seq peaks, MACS2 peaks, or HMMRATAC *center* and *nucleosome* states. **(C)** Distribution of Phastcons conservation score within vertebrates in 1000 bps window surrounding the summits of F-Seq, MACS2, and HMMRATAC calls.

Functional regulatory elements that harbor transcription factor binding sites are generally under higher evolutionary constraint (40). Success in transcription factor ChIP-seq data analysis is often benchmarked by high PhastCons score, a precompiled evolutionary conservation score at every base pair (41), at the peak center or summit, relative to the surrounding regions (42). We found that the summits of accessible regions detected by HMMRATAC in the GM12878 cell line also showed PhastCons enrichment in vertebrates (**Figure 3C**), indicating that these summits were likely locations of evolutionarily conserved regulatory elements and could represent functional regulatory elements.

### Method comparison with MACS2 and F-Seq

We next sought to compare the performance of HMMRATAC in identifying open chromatin regions to other popular methods for ATAC-seq. We chose MACS2 (11) and F-Seq (12) which had previously been shown to be the most accurate peak-callers in identifying open chromatin regions from DNase-seq (43). MACS2, originally designed for ChIP-seq, utilizes a local Poison model wherein enrichment is calculated relative to a local Poisson model. F-Seq invokes a Gaussian kernel-smoothing algorithm and then calculates enrichment based on the resulting Gaussian distribution. Although none of them utilize the intrinsic and unique features in ATAC-seq data we described before, MACS2 has been used to analyze ATAC-seq data in many published works (23,44–48), and MACS2 and F-Seq is part of the ENCODE analysis pipeline for ATAC-seq data (https://github.com/kundajelab/atac_dnase_pipelines). We didn’t include ZINBA, an algorithm based on the zero-inflated negative binomial model, in this manuscript, although it was used to analyze the GM12878 ATAC-seq dataset in Buenrostro et al.’s original work (7). ZINBA was reported as less sensitive than other methods while being applied to DNase-seq data (43) and we found the same conclusion (**Supplemental Figure S6**). Since this manuscript is not for reviewing all available methods, we decided to only show our improvement over the current best practices in the field. We tested the performance of the above algorithms using the two ATAC-seq datasets in the GM12878 cell line as the previous sections.

A major limitation in comparing the performance of identifying open chromatin regions is the absence of a valid “gold standard” (3). To overcome this problem, we chose to compare the performance of HMMRATAC and the other algorithms to three distinct independent datasets as the proxies of true and false set: 1) the active regions and heterochromatin from chromatin states analysis combining multiple independent histone features (32), 2) DNase-seq (5) and 3) FAIRE-seq (6) accessible regions of the same GM12878 cell line, originally called from traditional methods such as Hotspot (36) and F-Seq (12).

### Enrichment in chromatin states of active regions

The first proxy of the true set that we compared the algorithms to is referred to as “active regions.” These sites were defined through an integrative chromatin state segmentation using chromHMM (32,33) algorithm conducted by the ENCODE project. 8 histone modifications (H3K27me3, H3K36me3, H4K20me1, H3K4me1, H3K4me2, H3K4me3, H3K27ac, and H3K9ac), 1 transcription factor (CTCF) ChIP-seq and 1 whole genome input datasets in the GM12878 cell line were integrated through an unsupervised machine learning approach and each genomic location was annotated as one of 15 mutually-exclusive states. Two resulting states, “Active Promoters” and “Strong Enhancers”, were merged together to create our list of “active regions”. Additionally, the state annotated as “Heterochromatin” was used as a false set. In **Figure 2A** and **Supplemental Figure S7**, we showed some examples of consistent true positive calls from all algorithms and those false positive peaks called by MACS2 or F-Seq but not HMMRATAC, with default settings from each algorithm. A general observation is that conventional peak calling approaches can be misled by high ATAC-seq nucleosomal signals and call regions with no NFR signal as peaks. Full peak calls from default setting can be accessed through a UCSC genome browser track hub (http://biomisc.org/data/HMMRATAC_hub/hub.txt). However, the total number of such cases in the genome is strongly dependent on the cutoffs used by the individual algorithms. By tweaking cutoffs, false positives may disappear while sacrificing sensitivity. Therefore, we decided to test different cutoff values for each algorithm (see **Methods**). We found that HMMRATAC outperformed the other algorithms, in terms of precision, recall, and false positive rate in identifying these “active regions” while avoiding “heterochromatin” (**Figure 4** top row). The area under the curve (AUC) was then calculated for the recall vs. false positive rate curves (Area Under Curve of Receiver Operator Characteristics; AUCROC) and for the precision-recall curves (Area Under Precision-Recall Curve; AUPRC) (**Table 1**), after adding extreme points to the unreachable ends of the curves. HMMRATAC was proved to have better AUC than the two other methods. It worth noting that HMMRATAC calls larger peaks than the other two methods due to its merging of the adjacent nucleosome states to the center state. However, despite the larger average peak size, HMMRATAC identifies a comparable number of accessible regions in base pairs, as shown in **Supplemental Figure S8**, at comparable levels of sensitivity for “active regions”. This would also indicate that many peaks called by the other methods are falling outside of the “gold standard” regions, possibly contributing to their deficits in precision compared to HMMRATAC. Additionally, we retrieved 257 GM12878 super-enhancers from the dbSUPER (49) as a true set, assuming these regions are bound by an array of transcription factors and always been broadly accessible, and used the “heterochromatin” regions as false for an extra test. We found that HMMRATAC again performs better than other algorithms (**Supplemental Figure S9**). To study the effect of size-selection, we performed an *in silico* size-selection to restrict the fragments to those whose length was under 250 bps, then we tested the performance of HMMRATAC, MACS2, and F-Seq as shown before. We found that although the overall performance on the partial data decreases compared with full data set without size-selection, HMMRATAC can still outperform others (**Supplemental Figure S10**). Taken together, these results indicate that HMMRATAC is able to identify active chromatin regions with higher recall and precision and lower false positive rate, than MACS2 and F-Seq.

**Figure 4.**
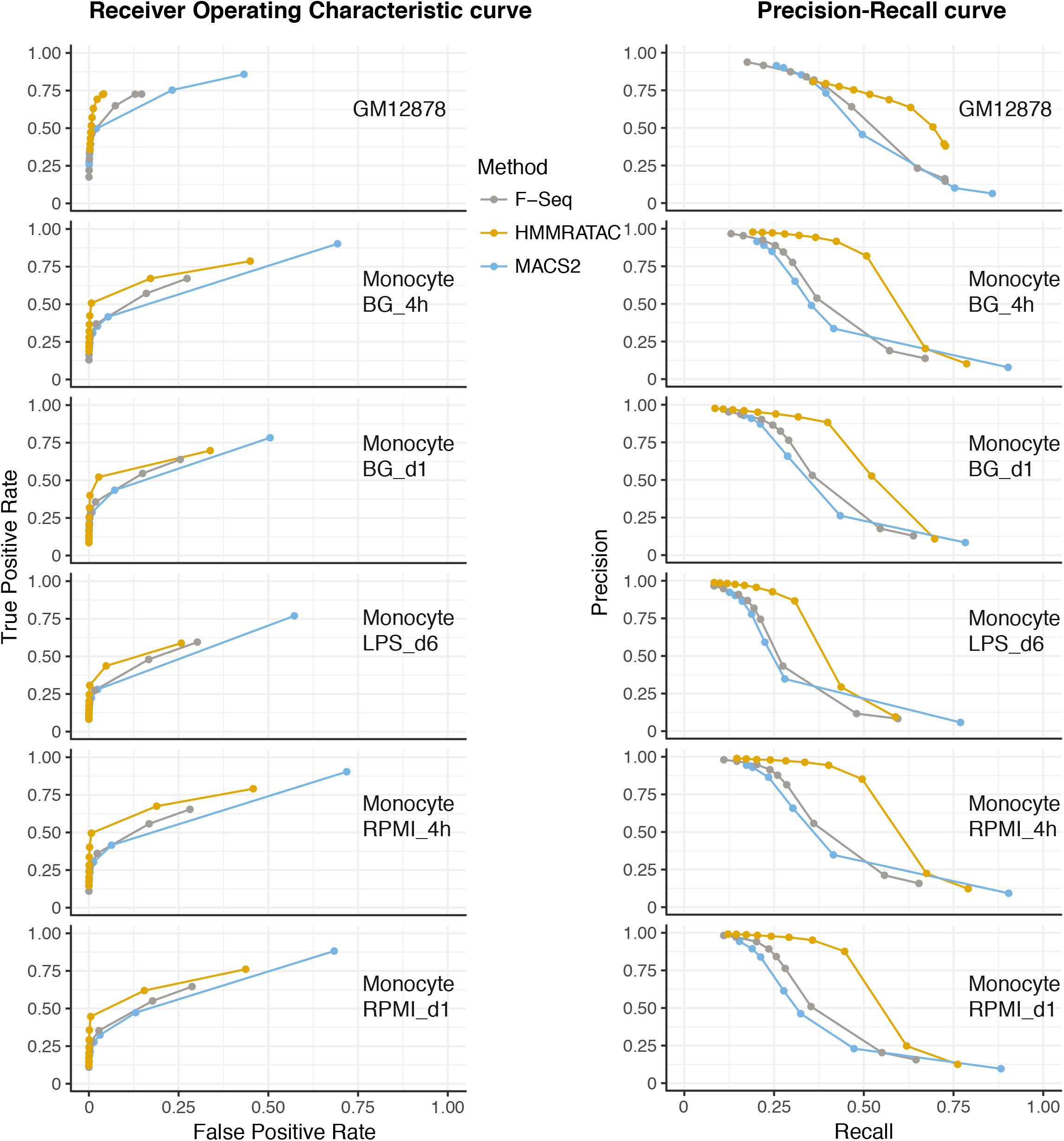
Comparison of performance between HMMRATAC, MACS2, and F-Seq. Left: Receiver Operator Characteristic (ROC) curves and right: precision-recall (PR) curves for the real positive and real negative pairs of 1. active chromatin states vs. heterochromatin for GM12878 cells, 2. active histone marks (either H3K4me1, H3K4me3 or H3K27ac) vs. heterochromatin (H3K9me3) for human monocytes (see **Methods** for detail).

**Table 1.**
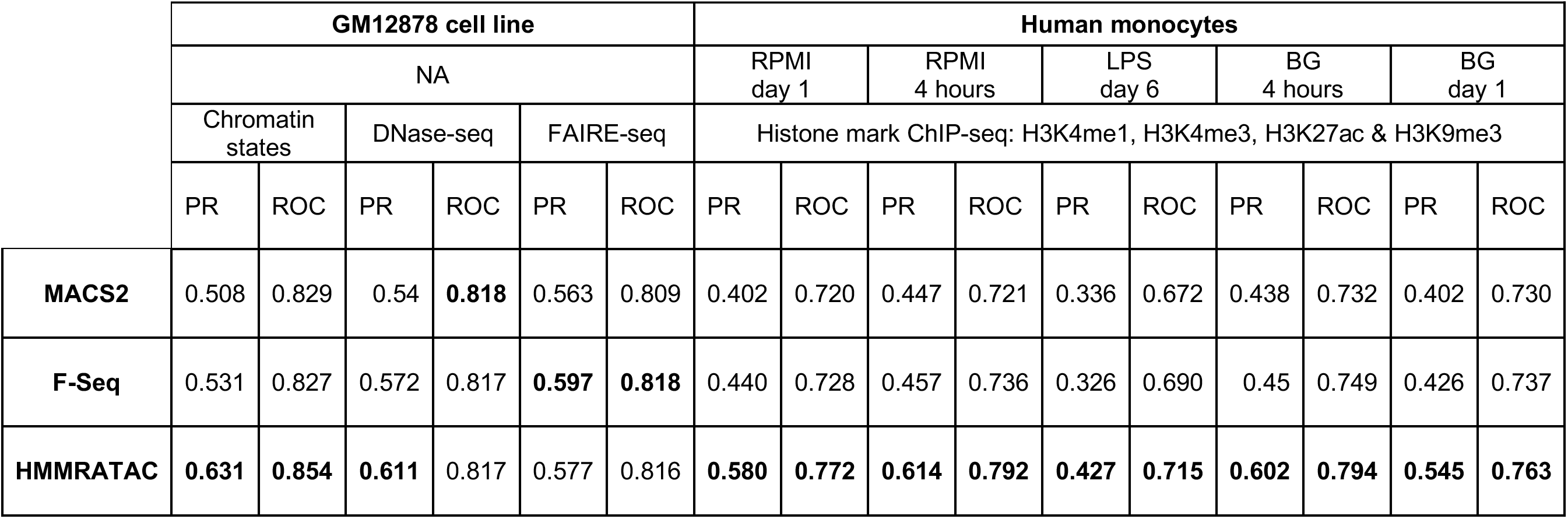
The Area Under the Curves (AUCs) of method comparisons of HMMRATAC, MACS2, and F-Seq. The AUC calculated from the Receiver Operator Characteristic (ROC) curves and Precision-Recall (PR) curves in **Figure 4** and **Supplemental Figure S11**. Because we can’t reach the extreme sides of the curves while tuning cutoff values, before we calculated AUC values, we added the following points to ROC curves– (0, 0) and (1, 1), and to PR curves – (0, 1) and (1, L_true_/(L_true_ + L_false_)). The L_true_ is the total length of the real positive set, and the L_false_ is the total length of the real negative set. The numbers of the best performance are shown in bold.

### Consistency with results from DNase-Seq and FAIRE-Seq

We then used the accessible chromatin regions detected by two independent assays DNase-seq and FAIRE-seq in GM12878 cell line as cross-validation of ATAC-seq analysis. We downloaded DNaseI hypersensitive sites (DHSs) (5) and FAIRE-seq (6) accessible regions called by ENCODE in the GM12878 cell line (35), then merged DHSs or FAIRE accessible regions that were within 2 kbps of each other to define our alternative true sets. We continued to use the heterochromatin states outlined above as our false set, although we eliminated any DHSs or FAIRE regions that overlapped a heterochromatin state region and *vice versa*. We found that HMMRATAC outperformed the other two methods in terms of AUPRC in recapitulating DHSs (**Supplemental Figure S11, Table 1**). MACS2 performed slightly better in AUCROC by 0.001. F-Seq was found to perform the best in terms of AUCROC and AUPRC with the FAIRE-seq data. Because the DNase-seq and FAIRE-seq regions were initially been called from traditional methods such as Hotspot (36) and F-Seq (12), the evaluations based on them are expected to be more biased to traditional approaches.

### Application for human monocytes

Thus far, we have shown the performance of HMMRATAC and other algorithms with data generated in the human GM12878 cell line. In order to determine our method’s performance with different data sets and different cell lines, we analyzed ATAC-seq data generated in human monocytes subjected to several chemical treatments and sequenced at different time points (23). In addition to ATAC-seq, Novakovic et al. performed ChIP-seq on the same cells, with the same treatments and time points, for the active histone marks histone H3K27ac, H3K4me1 and H3K4me3, and the repressive histone mark histone H3K9me3. We chose five separate data sets generated by that group: RPMI treatment after 4 and 24 hours, LPS treatment after 6 days and BG treatment after 4 and 24 hours. Using the active histone modification as the “Gold standard” and the repressive histone modification as the “real negatives” (See Method section for more details), we compared the performance of the algorithms in the same way we compared their performance in human data (**Figure 4** the second to the sixth row, **Table 1**). We found that HMMRATAC has superior precision, sensitivity, specificity, AUCROC and AUPRC over both MACS2 and F-Seq. Overall, this data indicates that HMMRATAC’s superior performance over the other algorithms is consistent across different data sets and different cell lines.

### Summary of method comparisons

The summary is shown in **Table 1**. We used a total of 16 independent measures in two different cell types and six different conditions to benchmark the performance of the three algorithms, including 8 AUCROC and 8 AUPRC. We found that HMMRATAC outperformed the other two methods in 6 of 8 AUCROC and 7 of 8 AUPRC. Furthermore, in all of the 12 evaluations of which the real positives were compiled by integrating multiple histone modification ChIP-seq datasets, HMMRATAC outperformed the other methods consistently. Overall, these results indicate that HMMRATAC is the most accurate and reliable peak-caller for ATAC-seq data analysis.

## DISCUSSION

HMMRATAC is designed to integrate the nucleosome-free and nucleosome-enriched signals derived from a single ATAC-seq dataset in order to identify open chromatin regions. For this reason, HMMRATAC should not be used on ATAC-seq datasets that have undergone either physical or computational size selection. For that same reason, single-end sequence libraries from ATAC-seq cannot be analyzed with HMMRATAC. However, we have shown that size selection results in a decrease in sensitivity and should not be used as a general practice in ATAC-seq protocols. Therefore, a reasonable way to evaluate the quality of ATAC-seq data for HMMRATAC analysis is to check the length distribution of the transposition fragments to ensure that large fragments are present and relatively abundant.

Identifying transcription factor footprints, wherein the DNA protected by a binding protein shows resistance to nonspecific DNA digestion, is an area of active research. Although footprint identification may be possible with certain datasets or for specific transcription factors, we believe that HMMRATAC identified summits, or the position with the most insertion events within the open region, are more robust to identify potential transcription factor binding sites. In fact, it has been previously shown that DNase-seq signal intensity at a binding motif is a better predictor of transcription factor binding than the strength of a footprint (50), and we believe this principle applies to ATAC-seq as well. However, a major reason for this result from He et al., was that many footprints were artifacts caused by the enzymes’ sequence bias. The Tn5 transposase also has a sequence bias, which would need to be corrected before an effective footprint analysis could occur. Several methods exist (51,52) that could correct the sequence bias from an ATAC-seq library. HMMRATAC could be used downstream of such sequence corrections, to reduce bias’ in peak calling, or upstream of the corrections, to identify the reads within peaks that need to be corrected for footprint or other downstream analysis.

It is possible to extend HMMRATAC to identify differentially accessible regions between two or more conditions. Most of the approaches for identifying such differences from ATAC-seq data would suffer from the same problems mentioned before, namely, that these methods only consider the relative strength of the signal and the differences in signal intensity between conditions. As we’ve shown here, the incorporation of the structure of regulatory chromatin into a model could increase the accuracy and sensitivity of calling accessible regions. We believe our strategy of “decomposition and integration” can be adopted in a differential accessibility analysis pipeline, by studying the differential NFRs and nucleosomal signals separately and then combining together.

ATAC-seq shows numerous advantages over other methods for identifying open chromatin regions, such as DNase-seq and FAIRE-seq, owing to its low starting material requirement and simple protocol. For these reasons, ATAC-seq has become one of the standard assays for locating regions of open chromatin, particularly as biomedical research continues to move toward translational research and precision medicine. Although the number of published ATAC-seq datasets increases rapidly, researchers are still using methods initially developed for ChIP-seq to analyze the data. We present HMMRATAC as a dedicated algorithm for the analysis of ATAC-seq data. We take advantage of the nature of the ATAC-seq experiment that due to the lack of size-selection while preparing the DNA library in its protocol, DNA fragments associated with well-positioned nucleosomes around open chromatin will be sequenced as well. Different from current methods where only the read enrichment is considered, HMMRATAC separates and integrates ATAC-seq data to create a statistical model that identifies the chromatin structure at accessible and active genomic regions. We have shown that HMMRATAC outperforms other computational methods in recapitulating open and active chromatin identified with other assays such as chromatin state analysis integrating multiple histone mark ChIP-seq, DNase-seq and FAIRE-seq. As HMMRATAC is a cross-platform and user-friendly algorithm, we envision it becoming the standard for ATAC-seq data analysis pipeline, replacing current methods designed for ChIP-seq analysis.

## DATA AVAILABILITY

The HMMRATAC source code, executable file, quick start guide, and javadocs can be downloaded from https://github.com/LiuLabUB/HMMRATAC. All peaks/regions called by HMMRATAC, F-Seq, and MACS2, as well as the various “gold standard” files, can be downloaded from the following URL: http://biomisc.org/download/HumanData.tar.gz.

## ACKNOWLEDGEMENTS

We thank Dr. Michael Buck, Dr. Marc Halfon, and Dr. Yijun Sun for their expertise and assistance throughout all aspects of our study.

## FUNDING

This work was supported by the startup fund from the University at Buffalo, the Buffalo Blue Sky Award from the University at Buffalo, the NIH/NCI Roswell Park CCSG grant (P30CA016056), and the NIH/NCI IOTN DMRC grant (U24CA232979).

## AUTHORS’ CONTRIBUTIONS

EDT and TL conceived and designed the computational method. EDT developed the software package, implemented the method and analyzed the data. EDT and TL wrote the paper. All authors read and approved the final manuscript.

## CONFLICT OF INTEREST

The authors declare that they have no conflict of interests.

