## Supplemental Figure S1-S11, Table S1-S4 for "HMMRATAC: a Hidden Markov ModeleR for ATAC-seq"

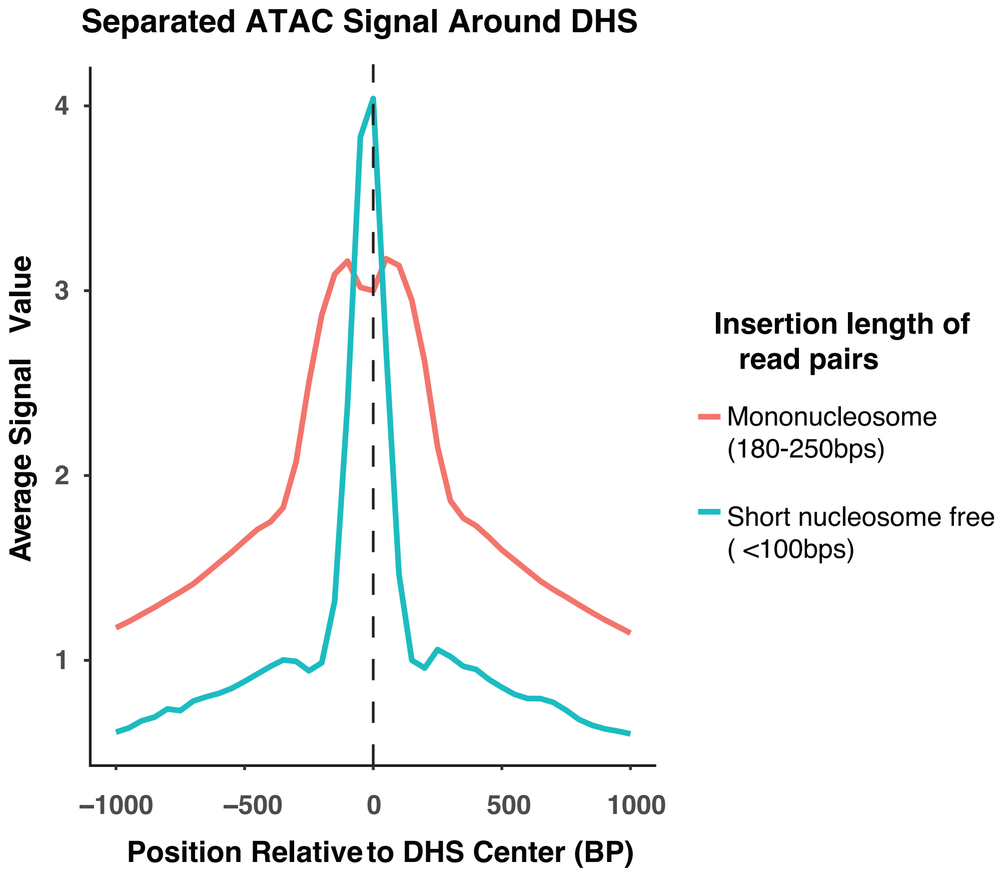

**Supplemantal Figure S1. Nucleosome-free and mono-nucleosome ATAC-seq fragments around DNaseI Hypersensitive sites.**

ATAC-seq signals, separated by fragment length, around DNase hypersensitive sites (DHS). Short nucleosome free reads are from read pairs with insertion length below 100 bps, mononucleosome reads are from those between 180 and 250 bps in length.

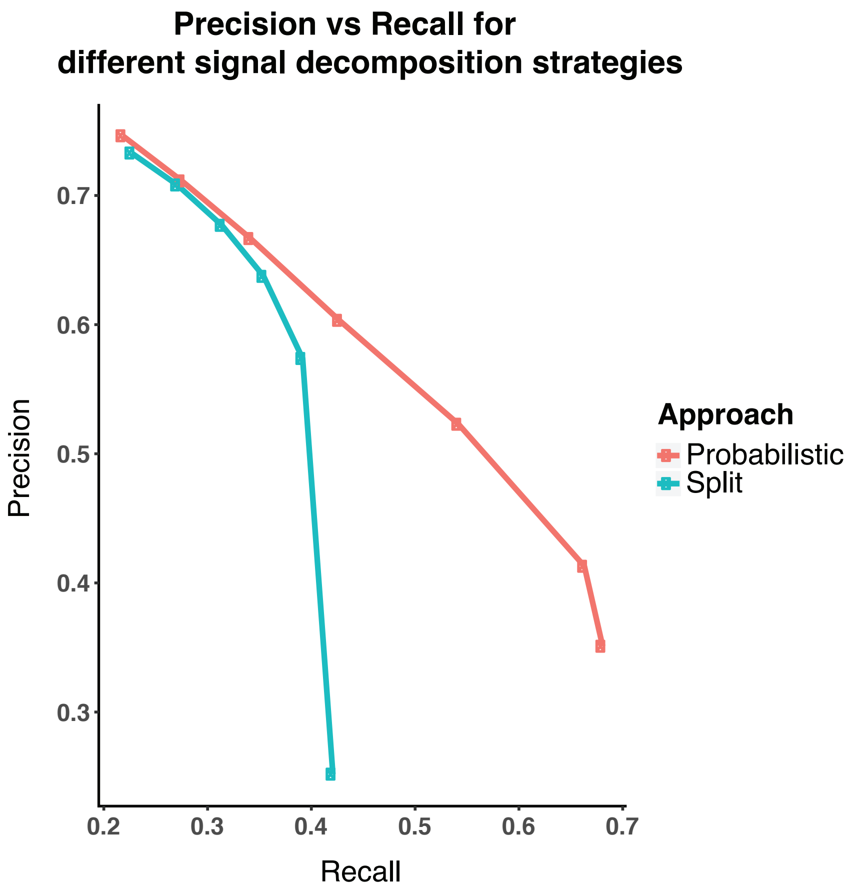

**Supplemental Figure S2. HMMRATAC probabilistic approach versus cutoff-based approach.**

Precision-Recall plots while using chromatin states as gold standard, comparing the probabilistic approach to signal decomposition used by HMMRATAC versus an alternative strategy where signal decomposition is accomplished using the cutoffs determined by Buenrostro et al.

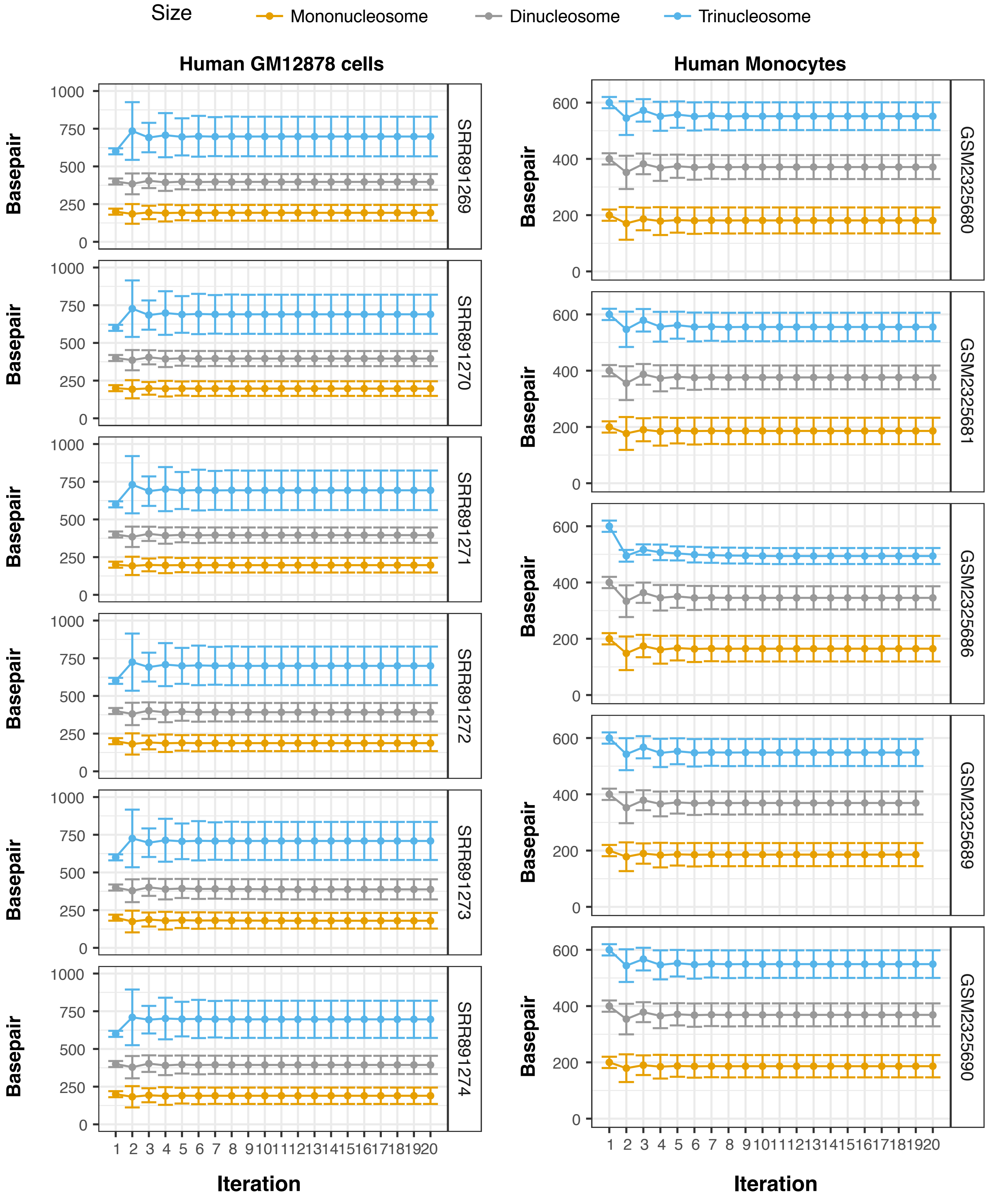

**Supplemental Figure S3. Mean and standard deviation parameters per EM iteration for three nucleosomal distributions, on human GM12878 and monocytes ATAC-seq data.** Note, the iteration #1 is in fact the initial parameters.

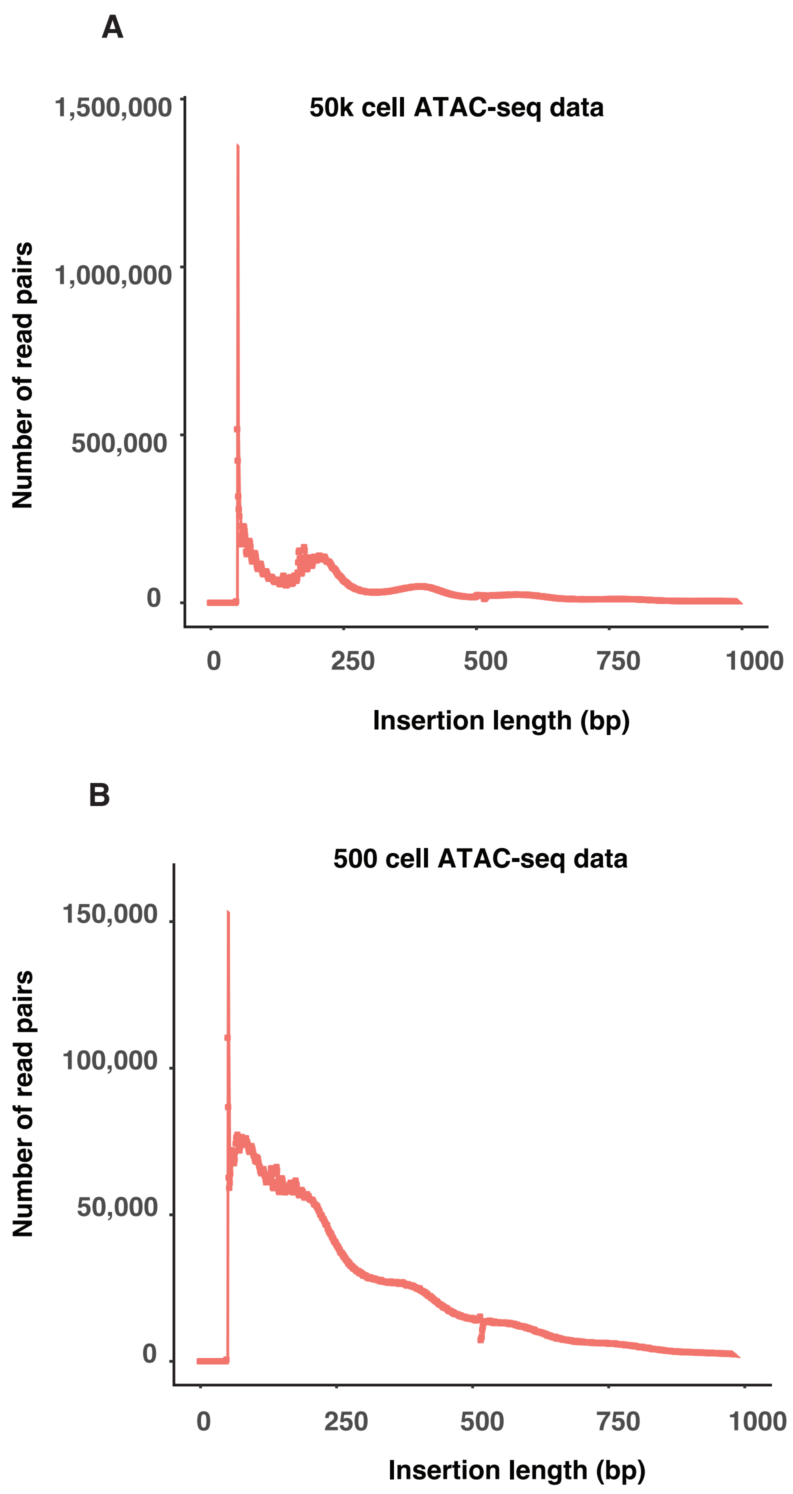

**Supplemental Figure S4. Fragment length distribution for 50,000 cells per replicate and 500 cells per replicate.**

Fragment length distributions were calculated from ATAC-seq assays conducted by Buenrostro et al. for 50,000 cells per replicate (**A**) and 500 cells per replicate (**B**).

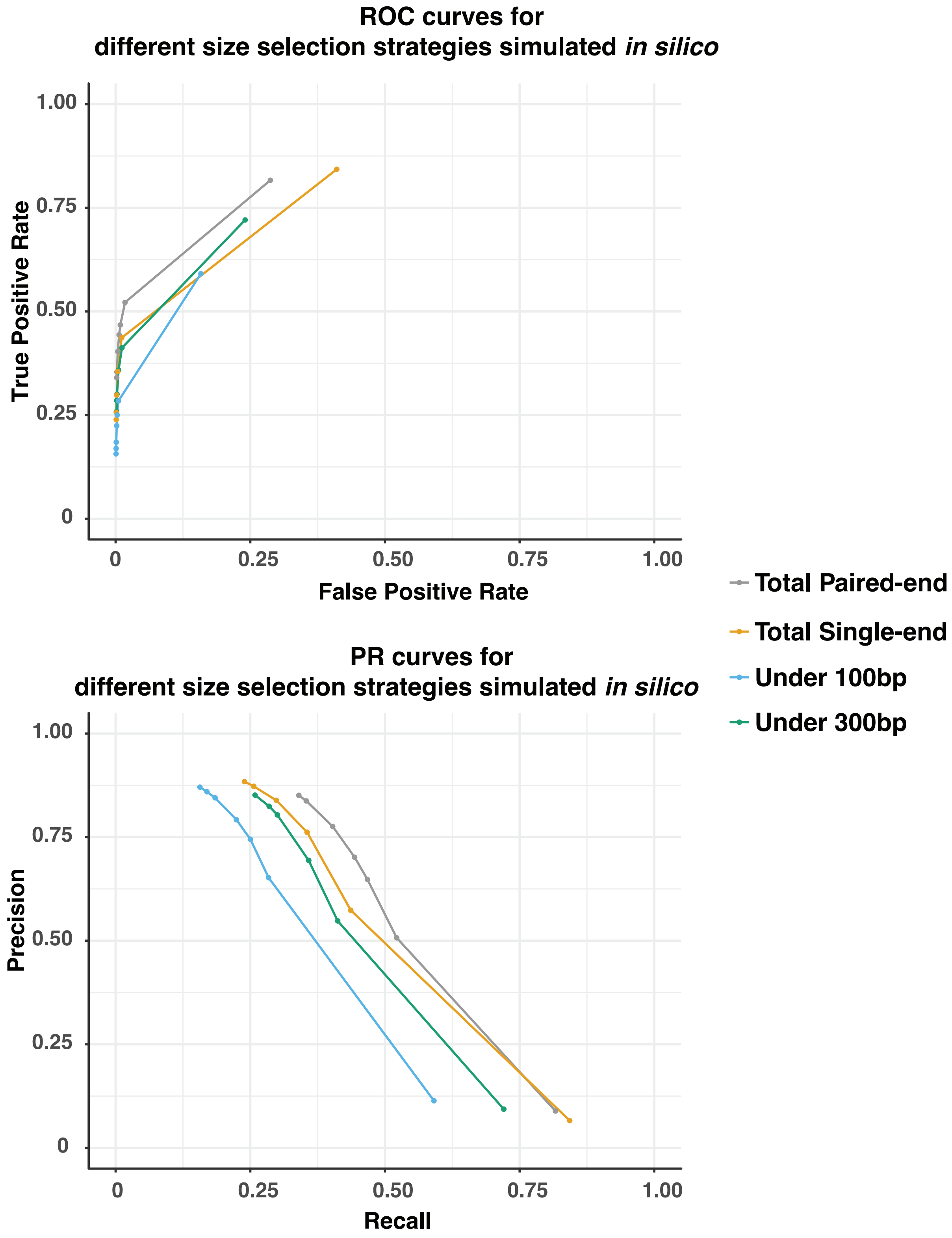

**Supplemental Figure S5. Effects under difference size-selection (*in silico simulation*) strategies.**

We used chromatin states in GM12878 as gold standard, comparing the different size-selection strategies by simulating Buenrostro et al. data *in silico*. We have four sets of data: 1) total reads with pairing information; 2) total reads without pairing information; 3) only read pairs with insertion size less than 100bp long; 4) only read pairs with insertion size less than 300bp long.

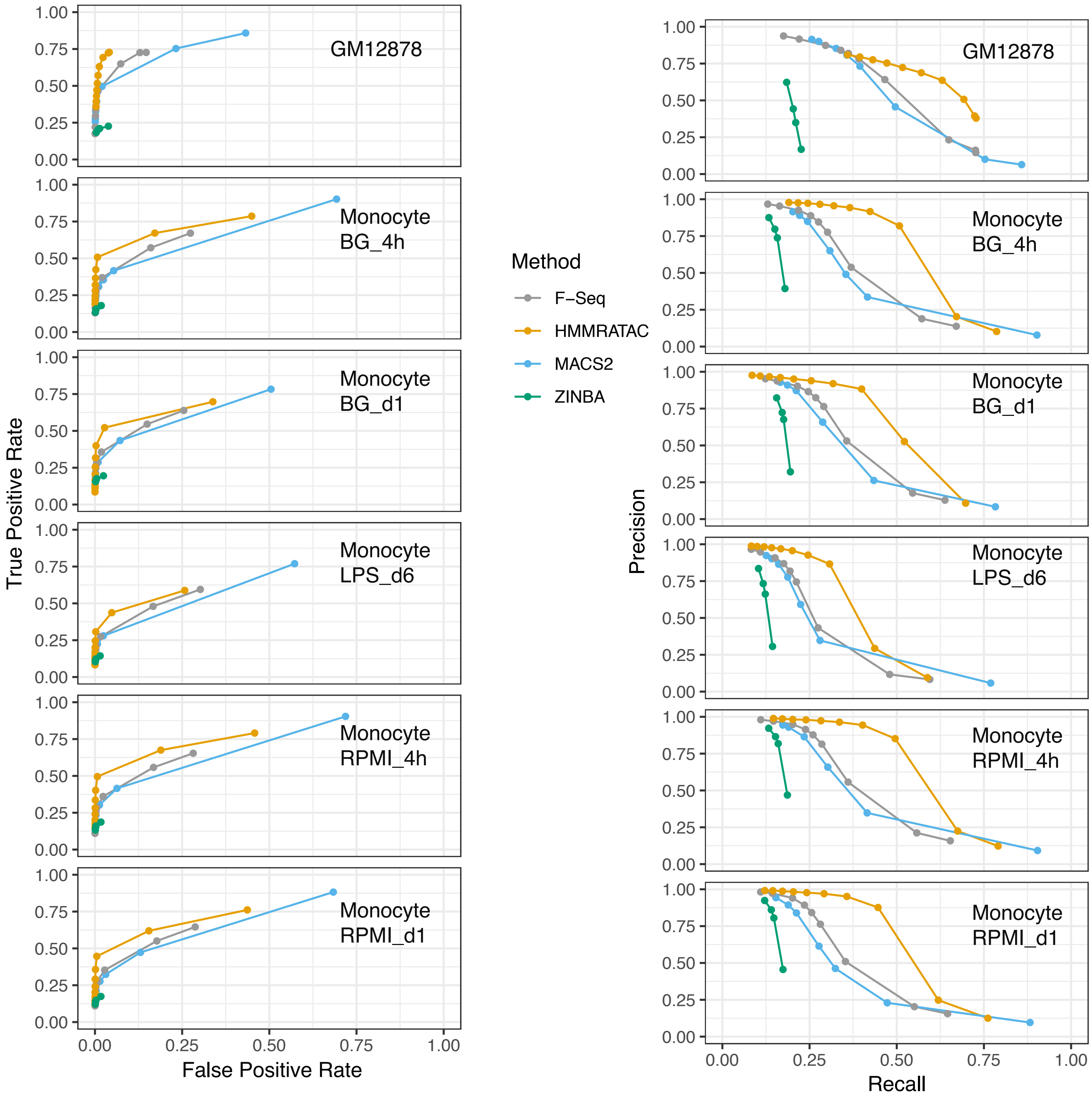

**Supplemental Figure S6. ROC curves and PR curves for testing HMMRATAC, F-Seq, MACS2 and ZINBA**

Same as the main Figure 4. Left: Receiver Operator Characteristic (ROC) curves and right: precision-recall (PR) curves for the real positive and real negative pairs of 1. active chromatin states vs. heterochromatin for GM12878 cells, 2. active histone marks (either H3K4me1, H3K4me3 or H3K27ac) vs. heterochromatin (H3K9me3) for human monocytes. Consistent with the comparison of peak callers for DNase-seq analysis (43), ZINBA underperforms other methods by a big margin. So we didn't include ZINBA in other analyses in this manuscript.

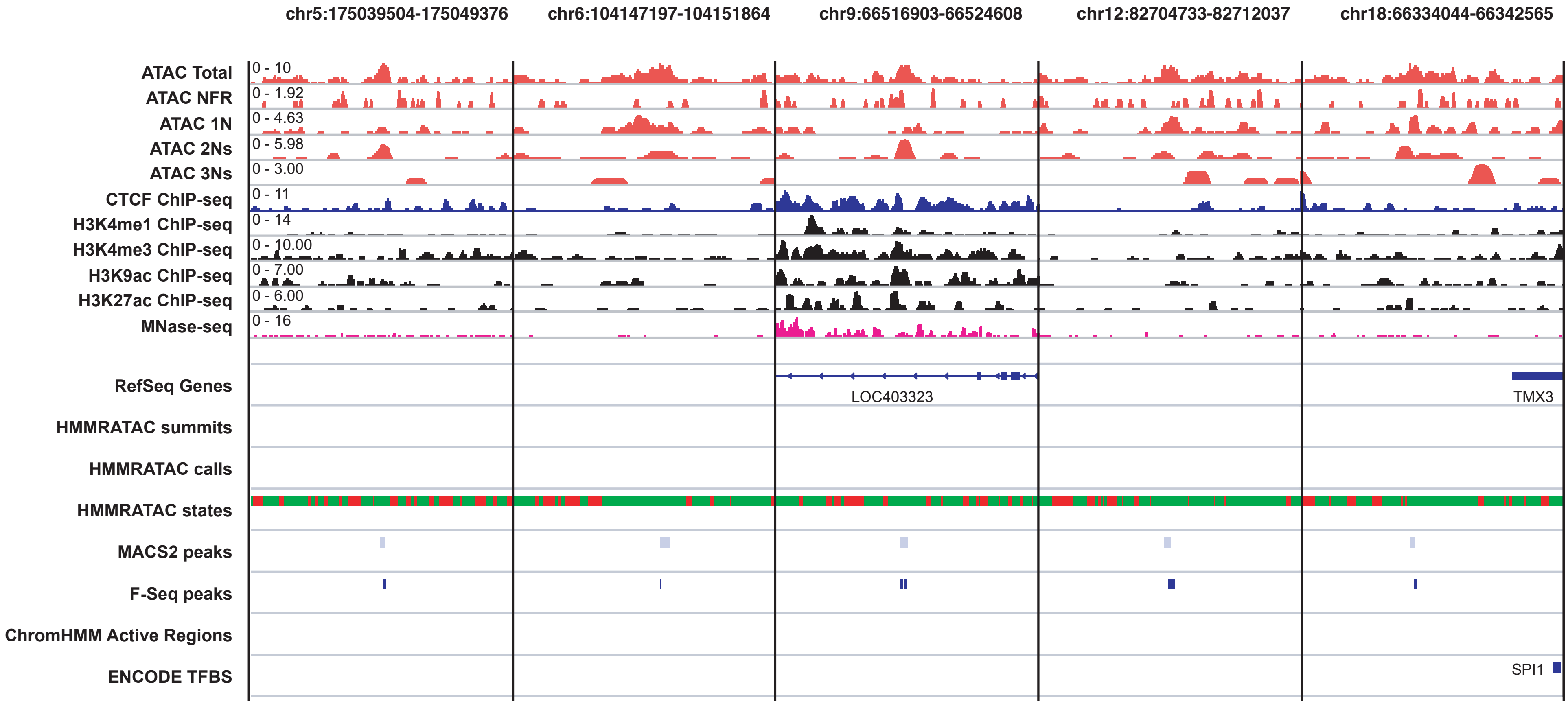

**Supplemental Figure S7. Some regions called by MACS2, F-Seq, but not by HMMRATAC**

We showed here five regions called by conventional peak callers but not by HMMRATAC, that all have a typical feature -- enriched with nucleosomal signals but low NFR signals. Similar to Fig 2, the green and red colored blocks in HMMRATAC state annotation track represent the nucleosome or background states.

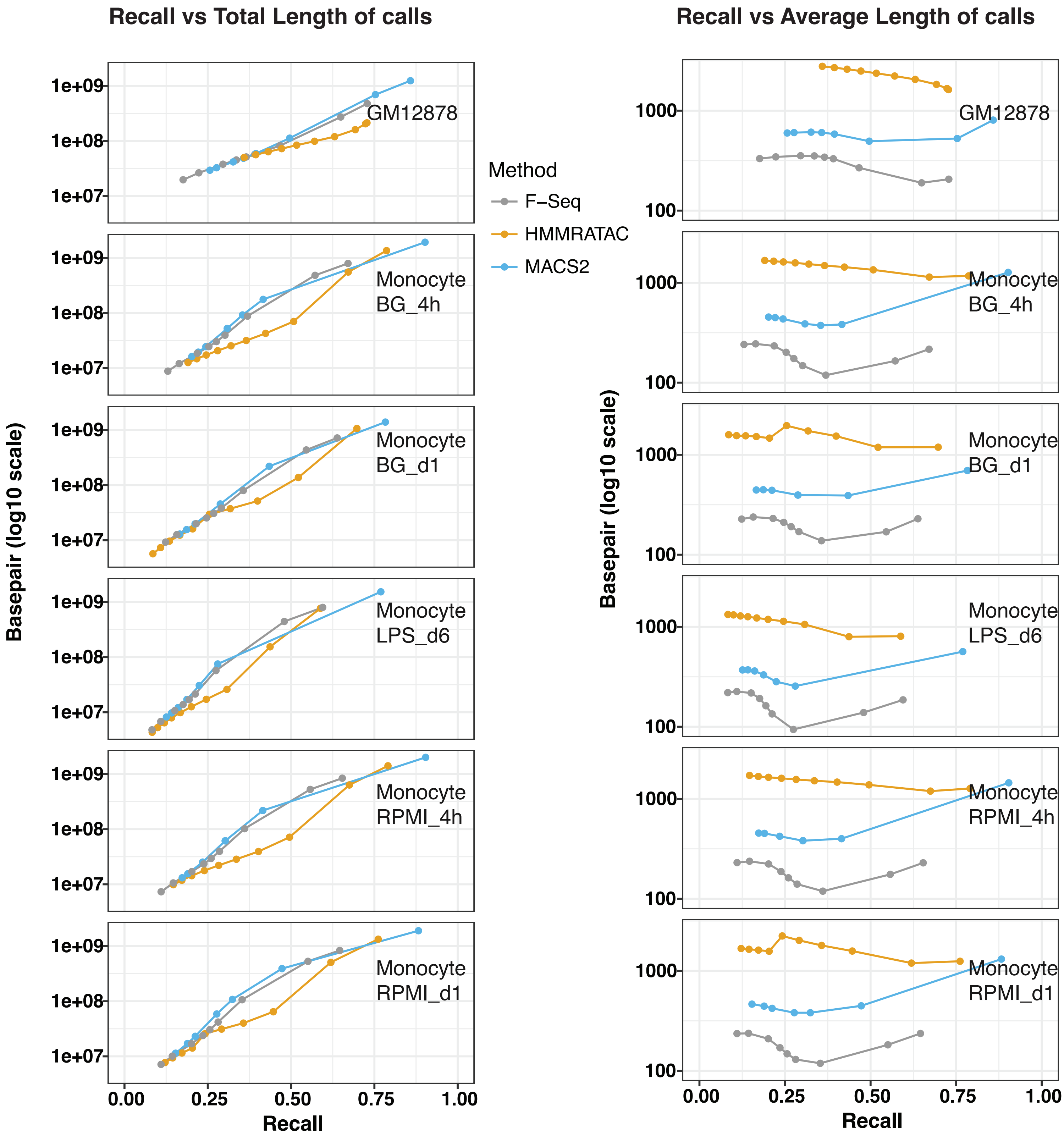

**Supplemental Figure S8. Recall/Sensitivity vs total length of calls or average length of calls from HMMRATAC, MACS2 and F-Seq**

The testing data, gold standard and real negatives are the same as Figure 4. Y-axis is in log10 scale.

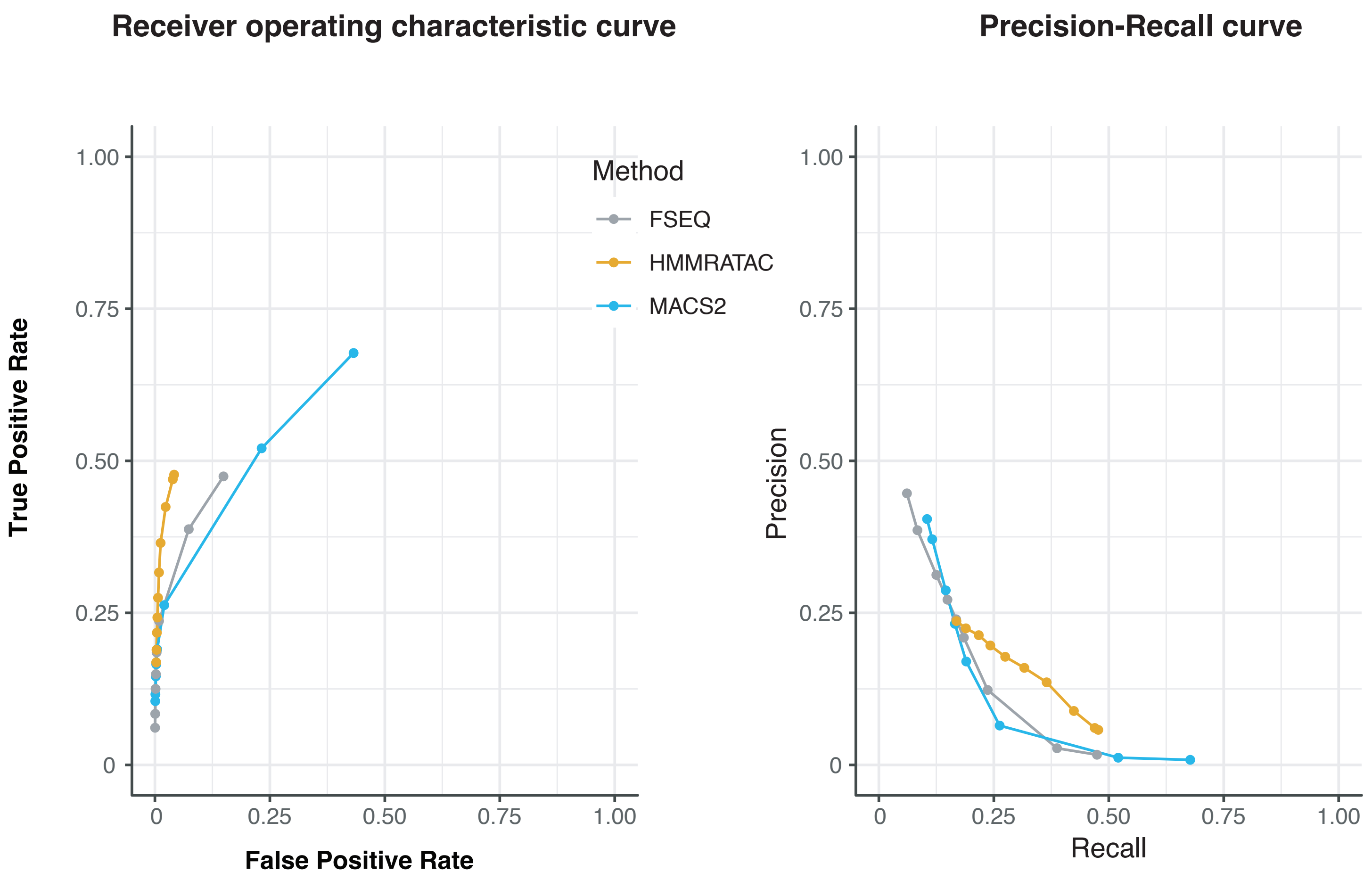

**Supplemental Figure S9. ROC curves and PR curves when testing super enhancers**

Evaluate methods on ATAC-seq data from GM12878 cell line while using super enhancers as true set, and heterochromatin chromatin states as false set.

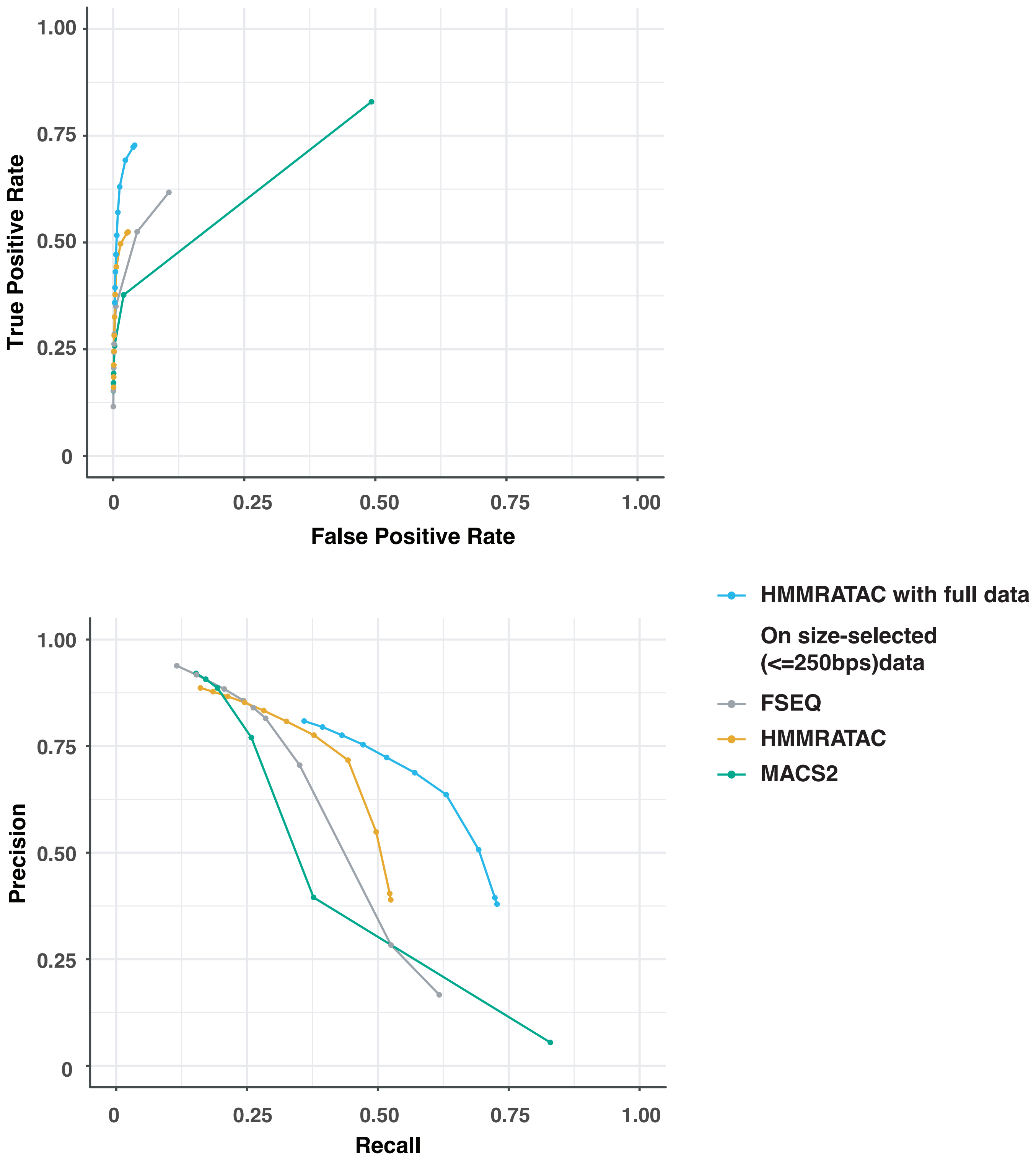

**Supplemental Figure S10. ROC curves and PR curves for testing HMMRATAC, F-Seq and MACS2 on size-selected data from GM12878 ATAC-seq**

We used chromatin states in GM12878 as gold standard as Figure 4, comparing the performance of different methods on the size-selected (<=250 bps) ATAC-seq data. Although the overall performance decreases compared with full dataset (HMMRATAC with full data vs HMMRATAC), HMMRATAC still outperforms other methods.

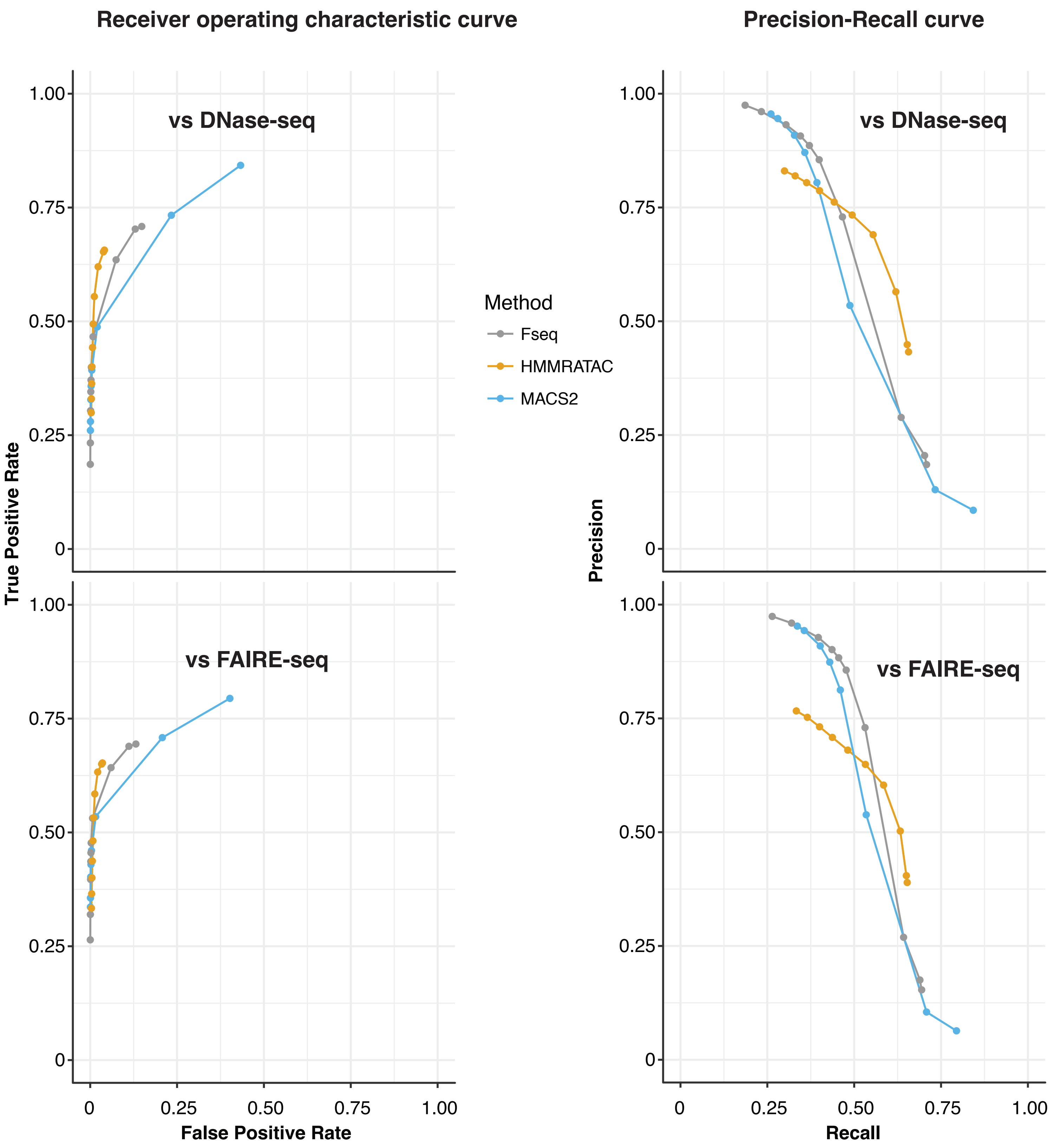

**Supplemental Figure S11. ROC curves and PR curves with DNase-seq or FAIRE-seq as standard**

Evaluate three methods on ATAC-seq data from GM12878 cell line. This figure shows ROC curvesand PRCs while using DNase-seq or FAIRE-seq peak calls from ENCODE as gold standard. The real negative set was defined as heterochromatin state from chromatin state analysis that do not overlap with any DNase-seq or FAIRE-seq peak calls, respectively.

| | | Transition parameters | | | Emission parameters | | | | |
|---|---|---|---|---|---|---|---|---|---|
| run | state | Background | Nucleosome | Center | NFR | 1N | 2Ns | 3Ns | |
| 1 | Background | 0.333 | 0.333 | 0.333 | 0.332 | 0.684 | 0.608 | 0.794 | from K-means |
| | Nucleosome | 0.333 | 0.333 | 0.333 | 0.863 | 1.731 | 1.913 | 2.574 | |
| | Center | 0.333 | 0.333 | 0.333 | 2.408 | 4.148 | 4.033 | 4.368 | |
| | Background | 0.951 | 0.049 | 0.001 | 0.064 | 0.26 | 0.541 | 0.773 | from Baum-Welch |
| | Nucleosome | 0.038 | 0.958 | 0.005 | 0.595 | 1.153 | 0.839 | 1.07 | |
| | Center | 0.001 | 0.009 | 0.99 | 1.527 | 2.75 | 2.856 | 3.436 | |
| 2 | Background | 0.333 | 0.333 | 0.333 | 0.332 | 0.684 | 0.607 | 0.794 | from K-means |
| | Nucleosome | 0.333 | 0.333 | 0.333 | 0.863 | 1.73 | 1.912 | 2.575 | |
| | Center | 0.333 | 0.333 | 0.333 | 2.409 | 4.148 | 4.031 | 4.37 | |
| | Background | 0.95 | 0.049 | 0.001 | 0.063 | 0.257 | 0.54 | 0.773 | from Baum-Welch |
| | Nucleosome | 0.038 | 0.958 | 0.005 | 0.594 | 1.152 | 0.838 | 1.07 | |
| | Center | 0.001 | 0.009 | 0.99 | 1.526 | 2.748 | 2.853 | 3.436 | |
| 3 | Background | 0.333 | 0.333 | 0.333 | 0.332 | 0.684 | 0.606 | 0.795 | from K-means |
| | Nucleosome | 0.333 | 0.333 | 0.333 | 0.863 | 1.73 | 1.909 | 2.577 | |
| | Center | 0.333 | 0.333 | 0.333 | 2.409 | 4.148 | 4.027 | 4.375 | |
| | Background | 0.95 | 0.049 | 0.001 | 0.063 | 0.257 | 0.538 | 0.774 | from Baum-Welch |
| | Nucleosome | 0.038 | 0.958 | 0.005 | 0.594 | 1.152 | 0.837 | 1.071 | |
| | Center | 0.001 | 0.009 | 0.99 | 1.526 | 2.748 | 2.85 | 3.439 | |

**Supplemental Table S1. Robustness to random initialization. We ran k-means clustering and HMM training three time with random initializations. We found the k-means results and the final HMM parameters are mostly identical among the three runs.**

| | Transition | | | Emission | | | | |
|---|---|---|---|---|---|---|---|---|
| | **1 background** | **2 nucleosome** | **3 center** | **NFR** | **N1** | **N2** | **N3** | **Sample** |
| **1 background** | 0.941646494 | 0.057626682 | 7.27E-04 | 0.00809468 | 0.00168701 | 0.26660351 | 0.51607081 | GM12878_50k |
| **2 nucleosome** | 0.022444289 | 0.973686683 | 0.00386903 | 0.44833745 | 0.91141612 | 0.75022441 | 0.71305135 | GM12878_50k |
| **3 center** | 8.97E-04 | 0.008881959 | 0.99022132 | 1.42875405 | 2.48954314 | 2.32517186 | 2.67086374 | GM12878_50k |
| **1 background** | 0.973592154 | 0.013258623 | 0.01314922 | 0.26534267 | 0.53730088 | 0.1012532 | 0.01111763 | GM12878_500 |
| **2 nucleosome** | 0.007559538 | 0.947031141 | 0.04540932 | 0.10623542 | 0.4278873 | 0.94871716 | 0.88893075 | GM12878_500 |
| **3 center** | 0.006251221 | 0.038300922 | 0.95544786 | 0.733387 | 1.29281661 | 1.18487558 | 1.08560785 | GM12878_500 |
| **1 background** | 0.94108237 | 0.049334524 | 0.00958311 | 6.44E-154 | 6.88E-155 | 6.92E-160 | 6.15E-165 | Monocyte_RPMI_d1 |
| **2 nucleosome** | 0.033945785 | 0.949682566 | 0.01637165 | 0.58035196 | 1.09937253 | 0.0198803 | 4.34E-06 | Monocyte_RPMI_d1 |
| **3 center** | 0.004682234 | 0.012768725 | 0.98254904 | 0.75578354 | 1.45120846 | 1.12300594 | 0.30983384 | Monocyte_RPMI_d1 |
| **1 background** | 0.942323253 | 0.050284474 | 0.00739227 | 0 | 0 | 0 | 0 | Monocyte_RPMI_4h |
| **2 nucleosome** | 0.036632826 | 0.948028679 | 0.01533849 | 0.53024286 | 1.05692383 | 0.0203743 | 3.60E-06 | Monocyte_RPMI_4h |
| **3 center** | 0.004923992 | 0.012448087 | 0.98262792 | 0.82656381 | 1.66069885 | 1.2126787 | 0.30101529 | Monocyte_RPMI_4h |
| **1 background** | 0.940676194 | 0.052896057 | 0.00642775 | 0 | 0 | 0 | 0 | Monocyte_LPS_d6 |
| **2 nucleosome** | 0.052891754 | 0.934903796 | 0.01220445 | 0.69024376 | 0.90409209 | 0.01901364 | 9.53E-11 | Monocyte_LPS_d6 |
| **3 center** | 0.007947584 | 0.015723289 | 0.97632913 | 0.73380067 | 1.23830927 | 0.95008077 | 0.20874454 | Monocyte_LPS_d6 |
| **1 background** | 0.965517163 | 0.028609696 | 0.00587314 | 0 | 0 | 0 | 0 | Monocyte_BG_d1 |
| **2 nucleosome** | 0.05141867 | 0.935152294 | 0.01342904 | 0.54936015 | 0.8994144 | 0.01724658 | 7.81E-06 | Monocyte_BG_d1 |
| **3 center** | 0.008998123 | 0.010309457 | 0.98069242 | 0.6529295 | 1.19554243 | 0.99172202 | 0.32536661 | Monocyte_BG_d1 |
| **1 background** | 0.935112863 | 0.0558214 | 0.00906574 | 0 | 0 | 0 | 0 | Monocyte_BG_4h |
| **2 nucleosome** | 0.043358826 | 0.940702059 | 0.01593911 | 0.64883274 | 0.98375378 | 0.01902861 | 4.94E-06 | Monocyte_BG_4h |
| **3 center** | 0.005043926 | 0.013647117 | 0.98130896 | 0.76665595 | 1.40078975 | 1.05988359 | 0.32775993 | Monocyte_BG_4h |

**Supplemental Table S2. HMM parameters from HMMRATAC Baum-Welch training on ATAC-seq data of GM12878 and monocytes.** The “NFR” column is for nucleosome free region signals, “1N” is for mono-nucleosome, “2Ns” is for di-nucleosome and “3Ns” is for tri-nucleosome signal.

| CistromeID | GEO Series | GEO Sample | Celltype | Tissue | Experiment Type | Condition |
|---|---|---|---|---|---|---|
| 65282 | GSE87218 | GSM2325690 | Monocyte | Blood | ATAC-seq | ATAC_RPMI_d1_8451 |
| 66431 | GSE85246 | GSM2262990 | Monocyte | Blood | H3K27ac | RPMI_d1_rep2_H3K27ac |
| 66436 | GSE85246 | GSM2262970 | Monocyte | Blood | H3K27ac | RPMI_d1_rep1_H3K27ac |
| 72242 | GSE85246 | GSM2263025 | Monocyte | Blood | H3K4me1 | RPMI_d1_rep2_H3K4me1 |
| 72245 | GSE85246 | GSM2263022 | Monocyte | Blood | H3K9me3 | RPMI_d1_H3K9me3 |
| 72249 | GSE85246 | GSM2263016 | Monocyte | Blood | H3K4me3 | RPMI_d1_rep1_H3K4me3 |
| 72259 | GSE85246 | GSM2263003 | Monocyte | Blood | H3K4me1 | RPMI_d1_rep1_H3K4me1 |
| 72277 | GSE85246 | GSM2262978 | Monocyte | Blood | H3K27me3 | RPMI_d1_H3K27me3 |
| 72281 | GSE85246 | GSM2262973 | Monocyte | Blood | H3K4me3 | RPMI_d1_rep2_H3K4me3 |
| 65283 | GSE87218 | GSM2325689 | Monocyte | Blood | ATAC-seq | ATAC_RPMI_4h_8448 |
| 66423 | GSE85246 | GSM2263019 | Monocyte | Blood | H3K27ac | RPMI_4h_rep1_H3K27ac |
| 66435 | GSE85246 | GSM2262977 | Monocyte | Blood | H3K27ac | RPMI_4h_rep2_H3K27ac |
| 72244 | GSE85246 | GSM2263023 | Monocyte | Blood | H3K4me1 | RPMI_4h_rep2_H3K4me1 |
| 72253 | GSE85246 | GSM2263012 | Monocyte | Blood | H3K9me3 | RPMI_4h_H3K9me3 |
| 72262 | GSE85246 | GSM2262998 | Monocyte | Blood | H3K4me3 | RPMI_4h_rep2_H3K4me3 |
| 72269 | GSE85246 | GSM2262989 | Monocyte | Blood | H3K27me3 | RPMI_4h_H3K27me3 |
| 72278 | GSE85246 | GSM2262976 | Monocyte | Blood | H3K4me3 | RPMI_4h_rep1_H3K4me3 |
| 72300 | GSE85246 | GSM2262948 | Monocyte | Blood | H3K4me1 | RPMI_4h_rep1_H3K4me1 |
| 65286 | GSE87218 | GSM2325686 | Monocyte | Blood | ATAC-seq | ATAC_LPS_d6_8455 |
| 66415 | GSE85246 | GSM2263053 | Monocyte | Blood | H3K27ac | LPS_d6_rep2_H3K27ac |
| 66440 | GSE85246 | GSM2262956 | Monocyte | Blood | H3K27ac | LPS_d6_rep1_H3K27ac |
| 72228 | GSE85246 | GSM2263046 | Monocyte | Blood | H3K4me3 | LPS_d6_rep2_H3K4me3 |
| 72232 | GSE85246 | GSM2263041 | Monocyte | Blood | H3K9me3 | LPS_d6_H3K9me3 |
| 72238 | GSE85246 | GSM2263031 | Monocyte | Blood | H3K4me3 | LPS_d6_rep1_H3K4me3 |
| 72257 | GSE85246 | GSM2263005 | Monocyte | Blood | H3K27me3 | LPS_d6_rep1_H3K27me3 |
| 72264 | GSE85246 | GSM2262996 | Monocyte | Blood | H3K4me1 | LPS_d6_rep1_H3K4me1 |
| 72287 | GSE85246 | GSM2262964 | Monocyte | Blood | H3K27me3 | LPS_d6_rep2_H3K27me3 |
| 72296 | GSE85246 | GSM2262953 | Monocyte | Blood | H3K4me1 | LPS_d6_rep2_H3K4me1 |

**Supplemental Table S3. The accession numbers for histone modification ChIP-seq datasets used to evaluate ATAC-seq analysis in human monocytes.**

| | | | | | | |
|---|---|---|---|---|---|---|
| 65291 | GSE87218 | GSM2325681 | Monocyte | Blood | ATAC-seq | ATAC_BG_d1_8453 |
| 66432 | GSE85246 | GSM2262988 | Monocyte | Blood | H3K27ac | BG_d1_rep1_H3K27ac |
| 66445 | GSE85246 | GSM2262937 | Monocyte | Blood | H3K27ac | BG_d1_rep2_H3K27ac |
| 72230 | GSE85246 | GSM2263043 | Monocyte | Blood | H3K4me3 | BG_d1_rep1_H3K4me3 |
| 72233 | GSE85246 | GSM2263040 | Monocyte | Blood | H3K9me3 | BG_d1_H3K9me3 |
| 72234 | GSE85246 | GSM2263039 | Monocyte | Blood | H3K4me3 | BG_d1_rep2_H3K4me3 |
| 72266 | GSE85246 | GSM2262994 | Monocyte | Blood | H3K4me1 | BG_d1_rep2_H3K4me1 |
| 72268 | GSE85246 | GSM2262991 | Monocyte | Blood | H3K4me1 | BG_d1_rep1_H3K4me1 |
| 72279 | GSE85246 | GSM2262975 | Monocyte | Blood | H3K27me3 | BG_d1_H3K27me3 |
| 65292 | GSE87218 | GSM2325680 | Monocyte | Blood | ATAC-seq | ATAC_BG_4h_8450 |
| 66443 | GSE85246 | GSM2262944 | Monocyte | Blood | H3K27ac | BG_4h_rep2_H3K27ac |
| 66444 | GSE85246 | GSM2262938 | Monocyte | Blood | H3K27ac | BG_4h_rep1_H3K27ac |
| 72229 | GSE85246 | GSM2263045 | Monocyte | Blood | H3K4me1 | BG_4h_rep1_H3K4me1 |
| 72241 | GSE85246 | GSM2263028 | Monocyte | Blood | H3K9me3 | BG_4h_H3K9me3 |
| 72248 | GSE85246 | GSM2263017 | Monocyte | Blood | H3K27me3 | BG_4h_H3K27me3 |
| 72252 | GSE85246 | GSM2263013 | Monocyte | Blood | H3K4me1 | BG_4h_rep2_H3K4me1 |
| 72283 | GSE85246 | GSM2262971 | Monocyte | Blood | H3K4me3 | BG_4h_rep2_H3K4me3 |
| 72313 | GSE85246 | GSM2262927 | Monocyte | Blood | H3K4me3 | BG_4h_rep1_H3K4me3 |

**Supplemental Table S3 (Continued). The accession numbers for histone modification ChIP-seq datasets used to evaluate ATAC-seq analysis in human monocytes.**

| State | Total | GC | H2az | H3K4me1 | H3K4me3 | H3K27ac | H3K27me3 |
|---|---|---|---|---|---|---|---|
| 1 background | 0.373 | 0.355 | 0.229 | 0.239 | 0.212 | 0.182 | 0.320 |
| 2 nucleosome | 0.588 | 0.415 | 0.529 | 0.562 | 0.504 | 0.476 | 0.639 |
| 3 center | 0.039 | 0.464 | 0.241 | 0.199 | 0.284 | 0.342 | 0.041 |

| State | H3K36me3 | H3K4me2 | H3K79me2 | H3K9ac | H3K9me3 | H4K20me1 |
|---|---|---|---|---|---|---|
| 1 background | 0.347 | 0.178 | 0.294 | 0.163 | 0.359 | 0.314 |
| 2 nucleosome | 0.584 | 0.490 | 0.557 | 0.455 | 0.579 | 0.627 |
| 3 center | 0.069 | 0.332 | 0.149 | 0.381 | 0.062 | 0.059 |

| State | ActivePromoters | Heterochrom | Insulator | PoisedPromoter | Repetitive | Repressed | StrongEnhancer |
|---|---|---|---|---|---|---|---|
| 1 background | 0.052 | 0.347 | 0.123 | 0.137 | 0.485 | 0.346 | 0.114 |
| 2 nucleosome | 0.167 | 0.636 | 0.487 | 0.424 | 0.489 | 0.628 | 0.422 |
| 3 center | 0.780 | 0.017 | 0.390 | 0.439 | 0.026 | 0.027 | 0.464 |

| State | Txn_elongation | Txn_transition | WeakEnhancer | WeakPromoter | WeakTxn |
|---|---|---|---|---|---|
| 1 background | 0.417 | 0.294 | 0.218 | 0.149 | 0.398 |
| 2 nucleosome | 0.579 | 0.657 | 0.590 | 0.463 | 0.590 |
| 3 center | 0.005 | 0.050 | 0.192 | 0.388 | 0.012 |

**Supplemental Table S4. Overlap between various chromatin features and HMMRATAC state annotations based on ATAC-Seq.**
